# Global health open-source goggles for fluorescence-guided surgery

**DOI:** 10.1101/2022.07.07.498317

**Authors:** Leonid Shmuylovich, Christine M. O’Brien, Karen Nwosu, Samuel Achilefu

## Abstract

Fluorescence-guided surgery (FGS), coupled with novel near infrared (NIR) fluorescent contrast agents, has significant potential to improve health but in current practice is less suitable for low resource settings. Although there are efforts to simplify FGS systems, technical, economic, and logistic challenges have hampered its global adoption. To overcome these impediments, we developed a low-cost, open-source, battery-powered and fully wearable FGS system called the fluorescence imaging augmented reality Raspberry Pi-based goggle system (FAR-Pi). Compared to current technologies that are expensive, bulky, and wall-powered, FAR-Pi has higher spatial resolution, depth of focus and fluorescence sensitivity. The FAR-Pi system has broad appeal by detecting the diverse fluorescence of NIR contrast agents undergoing clinical trials, as demonstrated by the successful identification of tumors in vivo with LS301, a tumor-targeting NIR contrast agent. As an open-source, inexpensive, and modifiable system, FAR-Pi promises to broaden access to FGS, thereby improving health worldwide.

## Introduction

Near infrared (NIR, 700-900 nm) fluorescence image-guided surgery (FGS) is a modality capable of high contrast delineation of important anatomy or diseased tissue in real time, at high spatial resolution, without using ionizing radiation, and at depths exceeding visible light imaging methods. These features set FGS apart as a versatile method for surgical interventions, and FGS has the potential to improve surgical outcomes for patients in both high-income countries and low-and-middle-income countries (LMICs). For example, in the setting of tissue conserving oncologic surgery, ensuring cancer-free margins during primary surgery reduces the need for costly re-excision (*1*). The burden of additional procedures is arguably even higher in LMICs where surgical resources are quite limited. FGS has the capacity to detect positive margins intraoperatively(*2, 3*), thereby ensuring that all malignant tissue is removed during the primary surgery and decreasing the rate of re-excision. Furthermore, FGS can enhance contrast of important anatomy such as nerves and blood vessels, thereby decreasing surgical complication rates. While patients in both LMICs and high-income countries would benefit from reduced surgical complication rates, mortality after cancer surgery has been shown to be disproportionately greater in LMICs (*4*) and therefore FGS-mediated reduction in surgical complication rate could have a profound impact on global cancer survival.

Due to its clear advantages, early clinical successes, and opportunities for broad impact (*2, 5*), there has been an explosion of new FGS hardware systems (*6*) and novel targeted contrast agents for a variety of surgical applications (*7, 8*). Current FGS hardware designs include standalone, handheld, and wearable systems, all of which have been tested in clinical trials (*6, 8*). However, as is the case for many medical innovations (*9*), the complexity and cost of existing FGS systems create barriers to clinical adoption.

To ensure that FGS is accessible globally, several factors must be considered. FGS systems must be designed with an understanding that the majority of health centers will not be able to reimburse the cost of expensive technology, and thus low-cost devices will be needed. Further, a compact form factor is necessary in small operating rooms that are unable to accommodate large devices. Integration of all components of the device into a single portable unit worn by the surgeon will eliminate the need for additional health professionals responsible for managing the device. This is particularly relevant as nearly all the FDA-approved FGS systems require an operator other than the surgeon to navigate the device, which imposes coordination efforts during surgery that are particularly challenging in medical centers that face surgeon and staff shortages. Finally, devices must be designed to withstand the variability inherent in clinical settings around the world. For example, battery-powered devices are preferred to prevent tethering surgeons to electrical cords and in settings where power is unavailable or outages may be frequent. In addition, they must be maintained without expensive contracts, custom parts, or complicated troubleshooting guides, flaws that have led the majority of “high end” medical equipment donations to sit broken or unused in low resource centers, especially LMIC hospitals (*10*).

Open-source hardware incorporating low-cost single board computers (SBC) like the Raspberry Pi SBC, microcontrollers like the Arduino, off-the-shelf inexpensive sensors and electronics components, and rapid prototyping tools like 3D printing and laser cutting present an opportunity for making medical devices more globally accessible (*11, 12*). The Raspberry Pi SBC in particular has been used to develop multiple low cost accessible medical and research devices including digital scanning microscopes (*13*), syringe pumps (*14*), ear-contactless auscultation of patients with COVID-19 (*15*), and high throughput COVID-19 serologic testing (*16*).

To make NIR FGS available globally, we have developed a fluorescence imaging augmented reality Raspberry Pi-based goggle system (FAR-Pi), which is an open-source hardware inspired redesign of our existing optical see-through goggle augmented imaging and navigation system (GAINS) (*17*). Our novel low-cost device is a fully wearable battery powered augmented reality system made of inexpensive off-the-shelf components including Raspberry Pi SBC and custom 3-D printed parts, that captures real time co-aligned visible and NIR image streams which can be directly superimposed over a surgeon’s field of view. To our knowledge this is the first Raspberry Pi-based system for simultaneous and coaligned visible and NIR imaging. By making the design open source, accessible, inexpensive, and modular, our system will support multiple clinical use cases, foster innovation, and democratize this previously inaccessible technology for the global betterment of human health.

## Results

The components of a dual visible and NIR fluorescence-guided surgery system can be broadly separated into distinct categories: 1) an NIR excitation light source, 2) optical filters for removing excitation light and separating visible from NIR light, 3) imaging sensors to detect visible and NIR fluorescent signals, 4) a computational device that controls all system components, processes incoming signals, applies relevant algorithms to make the signal clinically relevant, and outputs processed data, 5) a mechanism for displaying meaningful clinical output to users, and 6) a power source. Figure 1A demonstrates a block diagram of these general categories of FGS systems implemented in our previously published GAINS system(*17*), which excites an NIR fluorophore with a benchtop fiber coupled laser that is separate from the surgeon, detects coaligned and real time fluorescence and visible light from an area of interest with Aptina CMOS sensors embedded in customized circuit boards, processes this data with appropriate threshold algorithms, and presents this information to the user as an augmented reality projection coaligned with their field of view. To decrease the clinical footprint, complexity, and cost of existing FGS systems (*6*), we consider off-the-shelf open-source hardware inspired redesigns of each major FGS system component (Figure 1B), and combine these new designs to create the FAR-Pi system.

**Figure 1.**
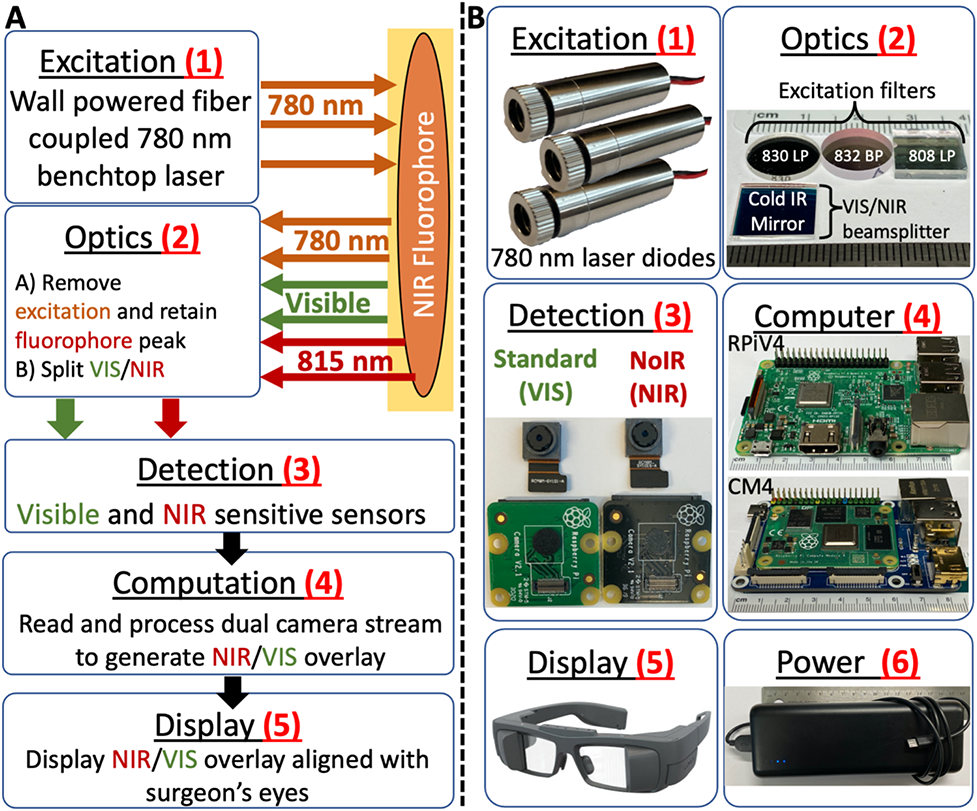
Summary of a representative fluorescence-guided surgery system and opportunities for simplification with an open-source low-cost off-the-shelf redesign. A) Block diagram of a representative see-through display fluorescence-guided surgical system which consists of a wall powered benchtop 780 nm laser module (1) that is capable of exciting NIR fluorophores, optical elements (2) designed to remove excitation light and split visible and NIR signals, VIS and NIR CMOS sensors with custom circuit boards and FPGA-based control module (3) designed to detect the signal, a laptop computer (4) that processes image sensor data to generate a false color NIR image, and an optical see-through display (5) that accepts input from the computer and provides the user with an NIR fluorescent signal superimposed over their field of view. The entire system is powered through wall power. B) To create a simpler, low-cost, and more accessible fluorescence-guided surgical system, the feasibility of replacing each existing component with less expensive off-the-shelf items was evaluated. Proposed alternatives include (1, Excitation): replacing the existing benchtop fiber-coupled laser module with an array of inexpensive 780 nm laser diodes, (2, Optics): using inexpensive cold-infrared mirror as a beamsplitter and a variety of potential excitation filters including an 830 nm longpass (LP) filter, an 832 nm bandpass (BP) filter, or an 808 nm LP filter, (3, Detection): detecting both visible and NIR signals using a Raspberry Pi v2 camera with a built-in NIR filter (left sensor) and an NIR-sensitive Raspberry Pi v2 camera without a filter (right sensor), (4, Computation): processing and outputting imaging data using a Raspberry Pi v4 (top) or Raspberry Pi Compute Module 4 (bottom) single-board computer, and (6, Power): powering the entire system using a rechargeable battery power bank rather than wall power.

### Near infrared and visible light detection

Commercial and preclinical NIR FGS systems achieve nanomolar fluorophore detection sensitivity by utilizing sensitive and often expensive CCD or CMOS detectors, in some cases coupled with custom electronics for image sensor processing(*6, 17–22*). An under-explored option for both fluorescent and visible light imaging that is less expensive and significantly simpler than existing FGS imaging sensors is the Raspberry Pi v2 camera module (RPiV2). The RPiV2 is available in a ‘standard’ NIR-insensitive model (with a built-in NIR filter), and in a ‘NoIR’ model with no NIR filter included. In addition, the RPiV2 supports miniaturization because the imaging sensor can be removed from the camera module circuit board and connected with flexible extension cables (Figure 2A). While the NoIR RPiV2 camera has been used to detect parathyroid autofluorescence (*23*), to our knowledge it has not been evaluated for use in a dual visible-NIR FGS system.

**Figure 2.**
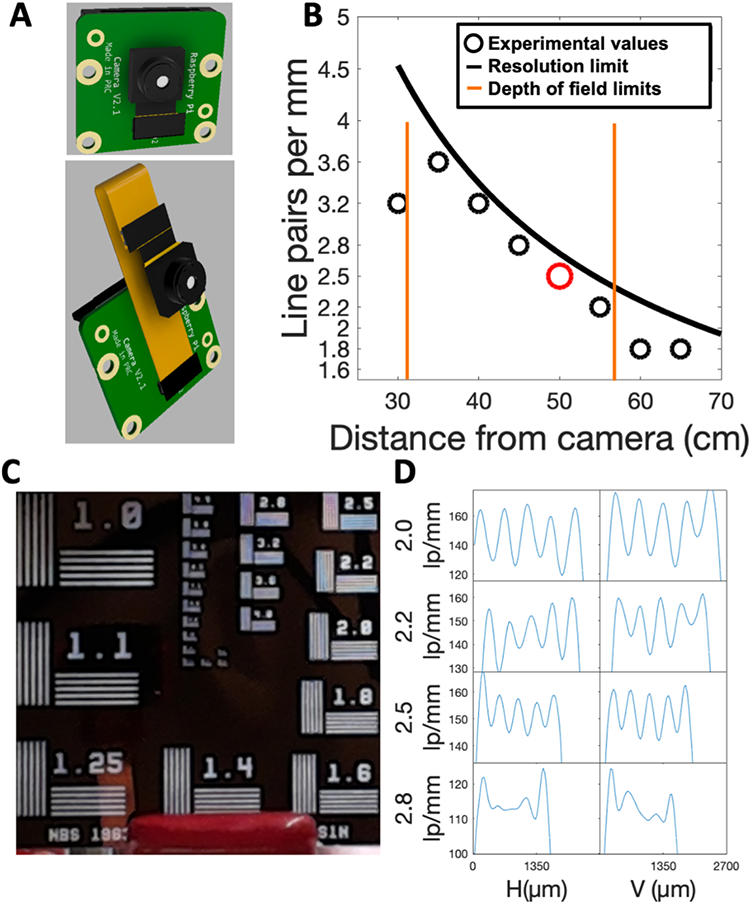
Raspberry Pi v2 camera has depth of focus and spatial resolution exceeding or matching existing fluorescence guided surgery systems. A) CAD model of the Raspberry Pi v2 camera module consisting of an IMX219 camera sensor and adjustable lens connected to a PCB. Miniaturization of imaging assembly is facilitated by disconnecting the camera sensor from the PCB and reconnecting it with a flexible extension cable. B) IMX219 sensor dimensions and lens field of view, focal length, and aperture can be used to predict camera spatial resolution and depth of focus limits as a function of object distance. Theoretical predictions were confirmed by determining the highest resolvable line pair group in a Thorlabs 1963A resolution test target at a given distance. Red circle indicates results at 50cm, which are shown in C) and D). C) Raspberry Pi v2 camera module-captured image of a Thorlabs 1963A resolution test target at a distance of 50 cm. D) Quantification of total pixel intensity across for horizontal and vertical line pairs within the 2.0, 2.2, 2.5, and 2.8 resolution target group numbers. Peaks correspond to centers of the white lines within each line pair, the highest group number with 5 discernible peaks (in this case group 2.5, corresponding to 200 µm resolution) defines the resolution limit.

Thus we tested whether RPiV2 standard and NoIR camera modules coupled with inexpensive optical components (Figure 1B ‘Optical Filtering’, ‘Detection’) and connected to a Raspberry Pi SBC (Figure 1B ‘Computation’) would support a simple, accessible, and low-cost implementation of FGS that matches the functionality of existing FGS systems.

#### Camera spatial resolution and depth of field characterization

To test the feasibility of the RPiV2 camera for FGS, first the spatial resolution and depth of focus limits of an RPiV2 camera were determined. Based on theoretical calculations (see Supplementary Methods, Section 2), an RPiV2 camera focused at 40 cm would have depth of focus limits between 31 and 56.2 cm. To test this prediction, the lens position was adjusted in an RPiV2 camera to achieve focus at 40 cm, and images of a standard spatial resolution test target were obtained at distances varying from 30 to 65 cm. The horizontal and vertical line pairs that could be resolved at each distance agreed with the theoretical predicted resolution limit when the distance was within the predicted in-focus range (Figure 2B, C, D).

These results demonstrate that an RPiV2 camera, which is the basis for visible and fluorescence detection in the FAR-Pi system, achieves horizontal and vertical spatial resolution ranging from 138 µm to 227 µm at distances between 35 cm and 65 cm. This resolution is in line with the 50-500 µm spatial resolutions reported in existing FGS systems (*6*) and represents an improvement upon the previously reported 320 µm spatial resolution using a video see-through GAINS system (*18*). The 25 cm depth of field far exceeds the 2-3 cm depth of field in both the GAINS system and commercial FGS systems (*6, 18*).

#### Coaligned visible and NIR imaging

Most FGS systems employ multiple cameras with optical filters and beamsplitter to achieve coaligned visible and NIR imaging (*6, 24*). A potentially simpler approach for real-time overlay of white light reflectance images with fluorescence images is using a single sensor coupled with a modified 4-channel RGB-IR Bayer filter (*6, 19, 25*). However, these off-the-shelf RGB-IR cameras such as the See3Cam_CU40 are not optimized for compactness and often go out of production, creating a fairly bulky (27cm^3^) unreliable system that is challenging to integrate into a wearable device. We postulated that 1) a cold infrared (IR) mirror could serve as an off-the-shelf and inexpensive beamsplitter, and 2) when coupled with a standard RPiV2 camera, could support coaligned real-time visible and NIR imaging. Figure 3A shows a to-scale schematic of the redesigned dual visible/NIR imaging assembly with a cold IR mirror operating as a visible/NIR beam splitter, while Figure 3B demonstrates a 3D cutaway illustrating the separate visible and NIR beam paths. Increasing the distance *ab* between camera and beamsplitter surface created sufficient clearance for thicker excitation filter designs (Figure S2) and thus facilitated the testing of multiple excitation filter combinations (filter optical densities in Figure 3A bottom panel).

**Figure 3.**
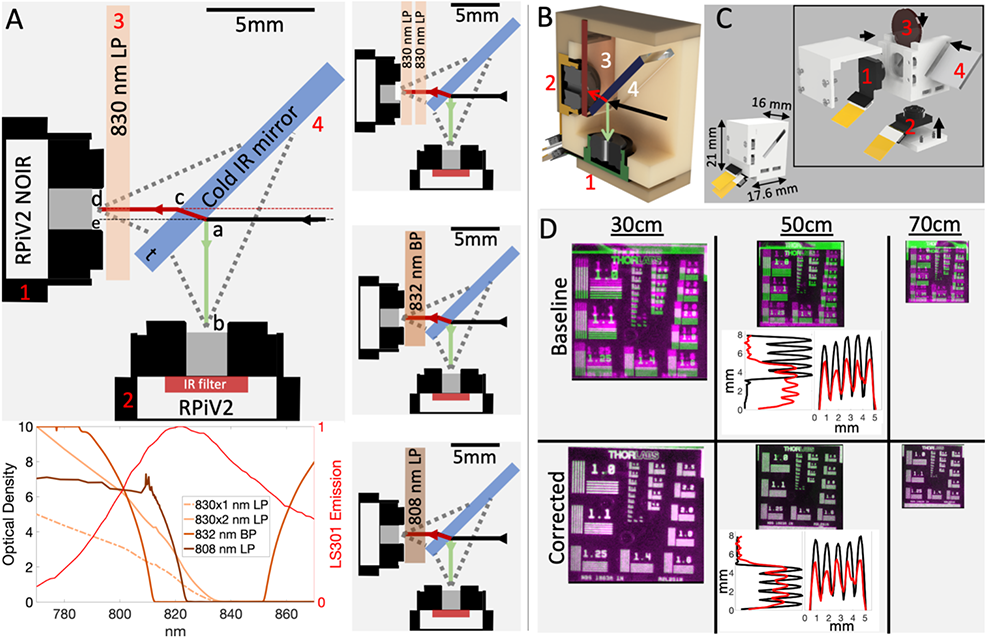
Raspberry PiV2 camera sensors can be combined with inexpensive optical components in a compact 3D-printed enclosure to achieve coaligned dual visible and NIR imaging. A) To-scale schematic demonstrating optically aligned cameras (items 1,2), optical components (items 3,4), and optical paths of visible (green line, path *ab*) and NIR light (red line, path *acd*). Majority of incident visible light is reflected off the cold IR mirror into an RPiV2 camera sensor (item 2) with internal IR filter. Majority of incident NIR light is transmitted and refracted through the cold IR mirror, and then passed through an excitation light filter before entering an RPiV2 camera NoIR sensor (item 1). Multiple excitation filter options were evaluated including a single 830 nm longpass (LP) Newport filter, two 830 nm LP filters placed in series, an 832 nm bandpass (BP) Edmund filter, and an 808 nm LP Semrock filter. In the lower panel of A), the optical density is plotted for each filter combination and compared to the normalized emission spectrum of LS301, a tumor-targeting NIR fluorophore(*26*). The cold IR mirror is positioned to ensure that field of view of each camera (dotted lines) does not extend beyond the edges of the mirror. B) Cutaway view of all components positioned within a 3D printed enclosure. C) Assembled enclosure and component-exploded view containing 3d printed enclosure, circular 12.7mm diameter longpass filter (item 3), square 12.7mm cold IR mirror (item 4), and RPiV2 NoIR (item 1) and standard IR-filter-containing (item 2) camera sensors. Note that the sensors are connected to flexible extension cables rather than connected directly to the RPiV2 PCBs. D) Superimposed visible and NIR images of a Thorlabs 1963A resolution test target captured by dual camera assembly at varying imaging distances. Inaccuracies in camera alignment result in a fixed vertical and horizontal pixel displacement that manifests as more prominent misalignment at larger distances. An affine transform defined from corresponding fiducial points obtained from the 50 cm NIR and visible images can be applied at any distance to produce near-perfect alignment between NIR and visible camera images. The middle panel shows horizontal and vertical line pairs from group 1.0 extracted from baseline (left) and affine-transform-corrected (right) resolution test target images acquired at 50 cm. Red peaks correspond to the line pairs extracted from the visible image, black peaks correspond to line pairs extracted from the NIR image. The degree of overlap between red and black peaks is a measure of coalignment between visible and NIR images in horizontal and vertical axes.

The relative positions of the NoIR and standard RPiV2 sensors in Figure 3A were calculated to ensure identical optical path and equal optical path length (*ab* = *ac+cd*). Custom 3D-printed components (Figure 3C and Figure S3) were designed to position and secure the RPiV2 NoiR and standard RPiV2 camera sensors as well as the square cold IR mirror and excitation filters in the 4 arrangements indicated in Figure 3A. After assembly of all components, the dual visible and NIR imaging assembly with a single 830 nm LP filter was a rectangular solid with dimensions of 21 mm x 17.625 mm x 16 mm (less than ¼ the volume of the single sensor RGB-IR See3Cam_CU40 camera), and weighed less than 5 g. Incorporating alternative excitation filters required minimal design adjustment (see Supplementary Methods, Section 5) without significant changes to the weight or size of the full assembly.

To test the degree of optical path alignment between the visible and NIR optical arms, images of a standard resolution test target at distances between 30 and 70 cm were obtained with both the NoIR and standard RPiV2 cameras and degree of image alignment was compared (Figure 3D, Baseline). When comparing NIR and visible images acquired from NoIR and standard RPiV2 sensors respectively, regardless of resolution target distance, horizontal alignment was within 1 pixel and vertical alignment was within 20 pixels. Despite this small deviation between NIR and visible sensor position, at a distance of 50 cm a 20 pixel difference results in a 3 mm vertical misalignment between the NoIR and visible images (Figure 3D, Baseline) and therefore a software based correction to ensure coalignment at every distance was developed. An affine transform was determined that co-aligned the 50 cm distance visible-sensor-derived image with the 50 cm distance NoIR-sensor-derived image. This same affine transform was applied to the visible-sensor-derived images acquired at 30 cm and 70 cm, and the transformed visible images aligned within 1 pixel of the distance-matched NoIR-sensor-derived images in both vertical and horizontal directions (Figure 3D, Corrected). Thus, in the FAR-Pi system a constant affine transform corrects the fixed physical offset between visible and NIR camera sensors and can be applied in real-time to visible images to ensure near-perfect alignment between visible and NIR images at any distance.

### Excitation light source

In fluorescence imaging, the excitation light source must have a spectral profile that overlaps with the excitation band of the desired fluorophore, and there must be a mechanism to maximize detection of fluorescence emission while minimizing detection of excitation light. For both indocyanine green (ICG) and a novel cancer-targeting NIR-fluorophore developed in our lab called LS301 (*26*), emission and excitation profiles are similar, with efficient excitation from 780-800 nm and peak emission at 815-830 nm (Figure 3A bottom panel).

Excitation at 780-800 nm can be achieved by use of broadband sources (halogen lamp or NIR light emitting diode (LED) array) coupled with appropriate optical filters, or with narrowband laser diodes (*6, 24*). However, incorporating an excitation light source into a wearable battery-powered ergonomic design is challenging. Existing FGS systems, for example, employ large cumbersome and expensive laser diodes that are fixed to a cart (*6, 17, 24*) and even if the light source is passed through optical fibers to a surgical headmount, the surgeon remains effectively tethered to the cart. Some preclinical FGS systems separate the surgeon from the excitation source completely – the imaging module is worn by the surgeon but the excitation source is placed on a tripod near the surgical field and separated from any wearable components (*18, 21*). Some groups have utilized less bulky and LED modules that may be wearable, but these LED modules require additional expensive optical filters and wall-powered non-wearable LED drivers (*21*). A halogen lamp (*22*) is a relatively inexpensive alternative, but also requires additional optical filters and poses a challenge with respect to being wearable given that it is wall-powered and generates significant heat.

To address these challenges, we evaluated whether an array of handheld 120 mW 3.3V 780 nm laser diode modules (Figure 1B, ‘Excitation’) could serve as a low-cost, wearable, and battery-powered alternative to existing bulky, non-wearable, and often expensive FGS excitation sources.

#### Characterization of low-cost alternatives to existing excitation light source

The 120 mW 780 nm laser diode modules are inexpensive (∼$18 each), readily sourced from commercial sources, compact (12 mm in diameter, 35 mm long), light weight (54 g) and operate at 3.3 V with a < 200 mA current draw. Measurement of the 780 nm laser diode’s spectral profile demonstrated a peak intensity at 783 nm with full width at half maximum of 1.5 nm (Figure 4A).

**Figure 4.**
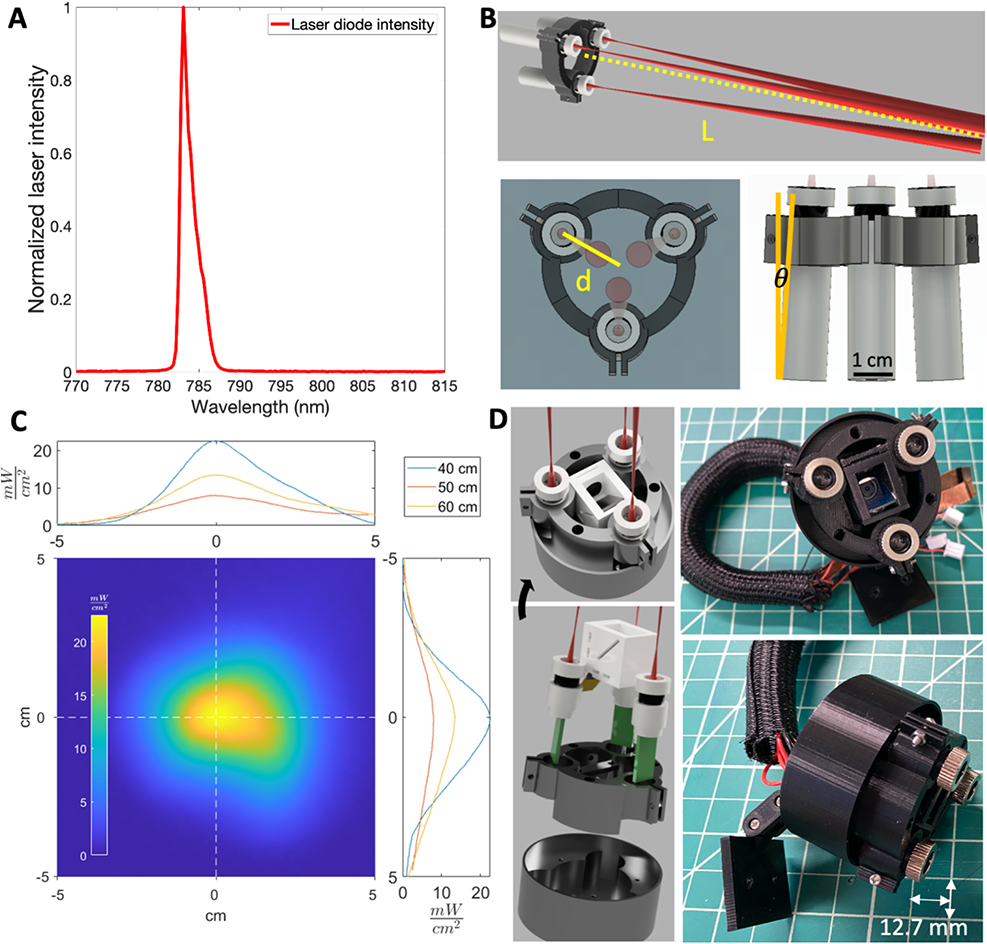
The FAR-Pi illumination module, consisting of a multi-purpose circular 780 nm laser diode array, delivers sufficient power for the FAR-Pi imaging module to operate at a working distance of 50 cm. A) Spectral characterization of a Laserlands 120 mW 780 nm laser diode demonstrating peak intensity at 783 nm with <2 nm full-width at half maximum. B) Parametric CAD model demonstrating a circular arrangement of laser diodes. The working distance *L* between laser array and the point where individual lasers intersect on the surgical field, and distance *d* between individual laser center and array center, determine the angle of laser inclination *θ*. C) Laser diode array spatial power distribution was measured at working distances of 40 cm, 50 cm, and 60 cm. The spatial power distribution at 50 cm is shown in the bottom left panel a 2 dimensional heatmap, and horizontal and vertical cross sections at 40, 50, and 60 cm are plotted in the top and right panels. At working distances of 50 cm, laser power exceeded 5 mW/cm^2^ over a 5.4 cm diameter area. D) CAD model (left) and physical component build (right) of a 3D-printed implementation of the illumination module where the FAR-Pi imaging module is positioned in the center of the laser diode array.

FGS applications typically utilize excitation power between 5 and 10 mW/cm^2^ (*6*). A battery powered circular array of three laser diodes was constructed (Figure 4B) and the spatial power distribution of the circular illumination area was determined at 40 cm, 50 cm, and 60 cm distances (Figure 4C). Maximum power was found to decrease from 20 to 5 mW/cm^2^ as the distance increased from 40 cm to 60 cm. At a distance of 50 cm, the irradiated power exceeded 5 mW/cm^2^ over a 5.4 cm diameter spot size.

The circular laser diode array provided a flexible baseline design that was utilized to position the VIS/NIR imaging module in the center of the laser diode array (Figure 4D). See Supplementary Methods, Section 6 for illumination module design and assembly details, including alternate designs that support additional excitation source optical filters and co-aligned white light sources. The full 3D printed assembly with laser diode array and centrally positioned dual-visible and NIR imaging module weighed 54 grams, and had cylindrical dimensions of 27 mm diameter base x 34 mm height (Figure 4D).

Thus a circular array of 3 compact laser diode modules is a sufficiently powerful, flexible, inexpensive, compact, wearable, and battery-powered replacement for bulky, wall-powered, and expensive excitation sources.

### Computation module integration

Existing FGS systems utilize personal computers such as Windows PCs or Linux-based mini-PCs to perform real time processing and subsequent display of imaging data(*6, 17, 18, 21*). While mini-PCs are smaller and less expensive than laptops, they cost at least $400, require wall-power, and are too bulky to be wearable. Using a single board computer (SBC) like the Raspberry Pi would provide a less expensive and wearable option, but limited SBC computational power may not support processing simultaneous camera streams in real time. We therefore tested if the most recent Raspberry Pi SBC, the Raspberry Pi V4 (RPiV4) as well as the Raspberry Pi Compute Module 4 (CM4), could read, process, and display simultaneous real-time camera streams and could control the laser diode array.

#### Dual Camera Streaming with the RPiV4

The RPiV2 NoIR camera circuit board was connected to the RPiV4 through the single camera serial interface (CSI) port, and the NoIR image stream was read and processed in real time using the python picamera package. The second RPiV2 camera in the FAR-Pi dual-camera enclosure was converted into a USB webcam with a Raspberry Pi Zero SBC running the ‘showmewebcam’ firmware (Figure 5A, B) and connected to the RPiV4 through a USB port. The RPiV4 successfully detected the USB camera, and OpenCV VideoCapture classes successfully read image frames from the USB camera. The picamera package was a more computationally efficient approach to capturing video frames than the OpenCV VideoCapture class. With the VideoCapture class we found it necessary to downscale image resolution in order to achieve full frame rate imaging. Using python threaded processes and OpenCV methods, images from both camera streams could be cropped, merged, transformed, and displayed to an external HDMI display or sent wirelessly to a device on a shared WiFi network in real time.

**Figure 5.**
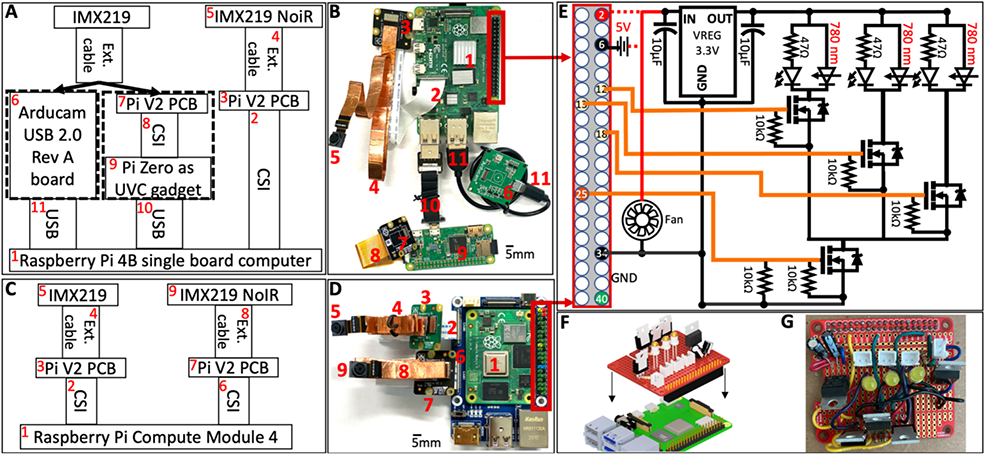
Raspberry Pi single board computers can simultaneously process two Raspberry Pi v2 camera module streams and control an external laser diode array. A) Schematic demonstrating methods for connecting two Raspberry Pi v2 (RPiV2) camera sensors (item 5 shows 1 camera, 2^nd^ camera sensor not connected) to a Raspberry Pi 4B (RPiV4) (item 1). RPiV2 sensors are connected to a data processing PCB with a sensor extension cable (item 4). One RPiV2 camera printed circuit board (PCB) module (item 3) can be connected directly with a CSI cable to the CSI port (item 2) of the RPiV4. Because the RPiV4 has only one CSI port, the second camera must first be converted into a USB-compatible camera which can then be connected via USB cable (item 10 or item 11) to one of four RPiV4 SBC USB-ports. Connecting the RPiV2 sensor to a custom 3^rd^ party PCB from Arducam (item 6) creates a USB-compatible camera. Alternatively, an RPiV2 camera PCB (item 7) can be connected to the CSI port of a Raspberry Pi Zero (item 9), and the Raspberry Pi Zero can function as a USB webcam by running the ‘showmewebcam’ firmware). B) The numbered items in schematic A) are implemented with physical components. C) Schematic demonstrating alternative implementation of A), where two RPiV2 camera sensors are connected directly to a Raspberry Pi Compute Module 4 SBC (RPiCM4). Because the RPiCM4 has two CSI ports, both cameras (items 5,9) can be connected directly to the RPiCM4 (item 1) without the need for additional hardware as in A). D) The schematic in C) is implemented with physical components. E) Schematic for custom circuit hardware connected directly to the 40 GPIO pins of either the RPiV4 or CM4. Power is drawn from either GPIO Pin 2 directly or from an external 5V supply. A 5V to 3.3V voltage regulator provides appropriate voltage to the laser diode array. N-channel MOSFET switches controlled by GPIO pins 12, 13, and 18 toggle individual laser diodes. An additional N-channel MOSFET switch toggles the entire laser diode array via GPIO pin 25. F) Schematic demonstrating how laser control circuitry is integrated with Raspberry Pi SBC GPIO pins. G) Assembled laser array control circuit ‘Hardware Attached on Top’ (HAT).

#### Dual Camera Streaming with the RPiCM4

The RPiCM4 (Figure 5C, D item 1) was coupled with a carrier board which exposed two CSI inputs to which standard and NoIR RPiV2 camera sensors were directly connected (Figure 5C, D). The python picamera package could detect, control, and read from both RPiV2 cameras directly without the need for computationally slower OpenCV VideoCapture classes.

#### Laser diode array control and synchronization with a Raspberry Pi Single Board Computer

We developed a custom ‘HAT’ (Hardware Attached on Top) circuit board that connects to a Raspberry Pi’s GPIO pins via a 40 pin female header and provides software-based control of each laser (Figure 5E-F). Specifically, this circuit provides a regulated 3.3V source for the laser diode array, powers a low current cooling fan to keep the RPiV4 from overheating, and connects 4 GPIO pins to 4 N-channel MOSFETS configured as switches. One switch (GPIO pin 25) is a master switch that toggles the entire laser array (on when GPIO pin is HIGH, off when the GPIO pin is low), while the remaining switches (GPIO pins 12, 13,18) toggle individual lasers. The HAT consists of readily available inexpensive off-the-shelf components like voltage regulators, transistors, capacitors, and resistors, and requires only basic hand-soldering skills to build.

Pulsed light imaging, where camera frame capture is synchronized with pulsed laser excitation so that laser excitation toggles between OFF and ON with consecutive frames, has been previously described to achieve real time NIR-background subtraction (*27*), though to our knowledge this has not been done using the Raspberry Pi SBCs or RPiV2 cameras. Using an ICG sample in a room with outside light coming through a window and overhead lights on, we tested whether the FAR-Pi system could support pulsed light NIR-background subtraction. We modified the device tree of both the RPiV4 and CM4 to allow GPIO pin 40 to serve as a readout for when the SBC starts and stops reading a frame (Figure 6A, camera pulse). Triggering the laser diode master switch to turn on with every other ‘frame end’ event and turn off after 70% of the duration of a single frame resulted in an image stream where Frames 1 and 3 had no laser excitation and no ICG fluorescence detected, while Frame 2 and 4 had laser excitation on and ICG fluorescence detected (Figure 6B). Longer laser excitation duration resulted in bleed through of laser excitation on consecutive frames. The lack of signal from a control vial of water next to the ICG vial confirmed that reflection of excitation light was not responsible for the detected signal. Background NIR light reflected off white paper was evident at the top of each frame regardless of whether laser excitation was present. The absolute value of subtracted consecutive frames (ΔF_n_ = |F_n_ – F_n-1_|) yielded a 30 fps video stream where background NIR signal was removed while excited fluorescence signal was preserved for each calculated frame (Figure 6C).

**Figure 6.**
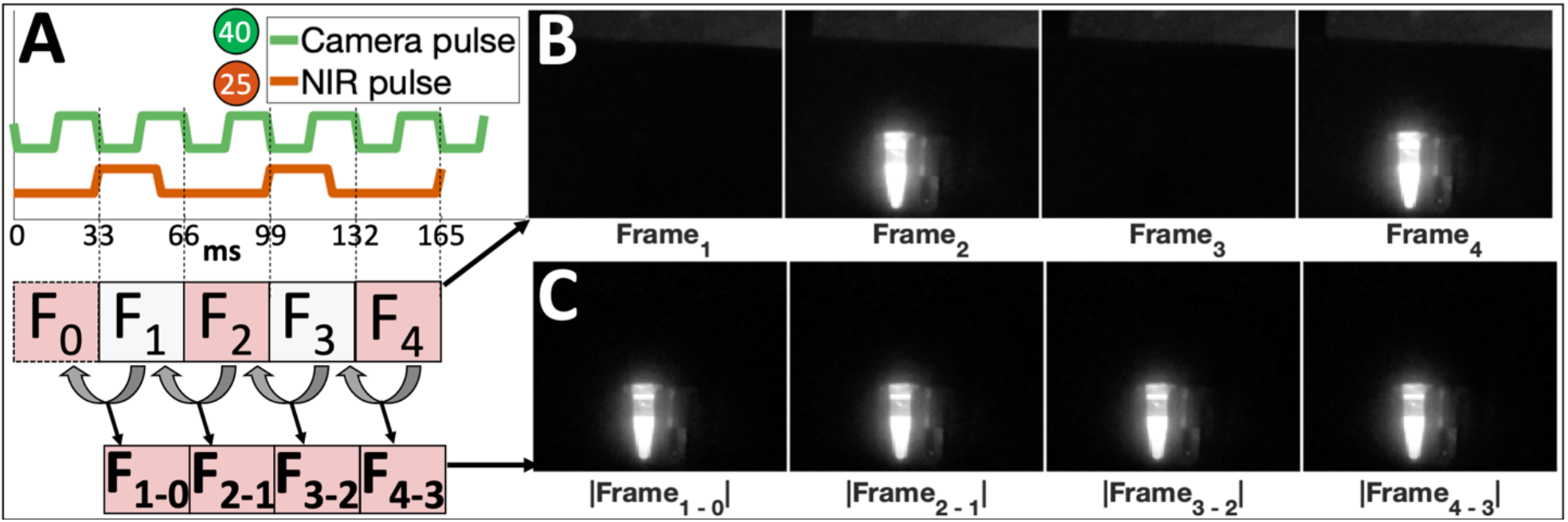
Synchronization of laser excitation with alternating camera frames facilitates real-time NIR-background subtraction. A) The device tree of the Raspberry Pi computer is modified to toggle GPIO pin 40 high/low with camera frame read start/stop events, generating a square wave (camera pulse, green) with 33 ms period when imaging at 30 frames per second. B) Triggering GPIO pin 25 at the end of every other frame (double the period of the camera pulse) toggles laser excitation with each frame, resulting in detected fluorescence from an Eppendorf tube filled with indocyanine green (ICG) on frames 2 and 4 in a sequence of 4 consecutive frames acquired with a NoIR RPiV2 camera fitted with a Newport 830 nm longpass filter. Note background NIR signal from room lights reflected off white paper near top of each acquired frame. C) The absolute value of the difference between the current frame and the preceding frame generates a real time video stream preserving the emitted fluorescence signal and removing the background NIR signal.

Thus the RPiV4 and CM4, when coupled with RPiV2 cameras, laser diodes, and the appropriate laser diode control circuitry, both have sufficient processing power to provide a compact and wearable FGS computational module capable of dual visible/NIR imaging with pulsed laser excitation and real time NIR-background-subtracted video streaming.

### Integrating imaging and illumination modules to align camera and surgeon’s view

A fundamental challenge in see-through FGS systems is ensuring coalignment between the fluorescence signal that is projected onto the see-through heads up display (HUD) and the surgeon’s real world view. Specifically, when the FAR-Pi system detects the fluorescent glow of a tumor-targeting NIR fluorophore, that signal does not necessarily carry any anatomical landmarks. For this signal to be useful to the surgeon when displayed on a see-through HUD, the position of the projected NIR signal must precisely overlap with the position of the tumor through their vision, and any misalignment could result in the surgeon misinterpreting where in the surgical field the fluorescent signal is emanating from. The degree of misalignment depends on the position of the FAR-Pi imaging module relative to the surgeon’s eye.

We identified 3 alternative positions for the FAR-Pi imaging module and for each alternative assessed the degree of misalignment between the FAR-Pi camera view and surgeon’s view and evaluated software-based strategies for image re-alignment. In the first design the FAR-Pi imaging module is placed in the center of the illumination module laser array (Figure 7A). This design ensures that the center of laser illumination is always at the center of the camera FOV, but there is significant horizontal and vertical disparity between the camera light path and eye light path. In the second design, the FAR-Pi imaging module was decoupled from the center of the illumination module and centered at eye level between the eyes (Figure 7B, left panel). This position eliminates vertical disparity and reduces horizontal disparity between the camera and eye light paths. In both of these designs, images obtained at distance of 50 cm by the FAR-Pi imaging module (Figure 7A, B ‘RPiV2 view’) and surrogate eye cameras (Figure 7A, B ‘eye view’) were not aligned, resulting in significant misalignment when overlaying the FAR-Pi captured image with the surgeon’s view (Figure 7A, B ‘Raw Overlay’). In both cases these images could be used to calibrate a 4-point projective transformation, and applying this transform to the FAR-Pi imaging module images corrected the image misalignment and allowed for error-free image overlay in both designs (Figure 7A,B, ‘Corrected’).

**Figure 7.**
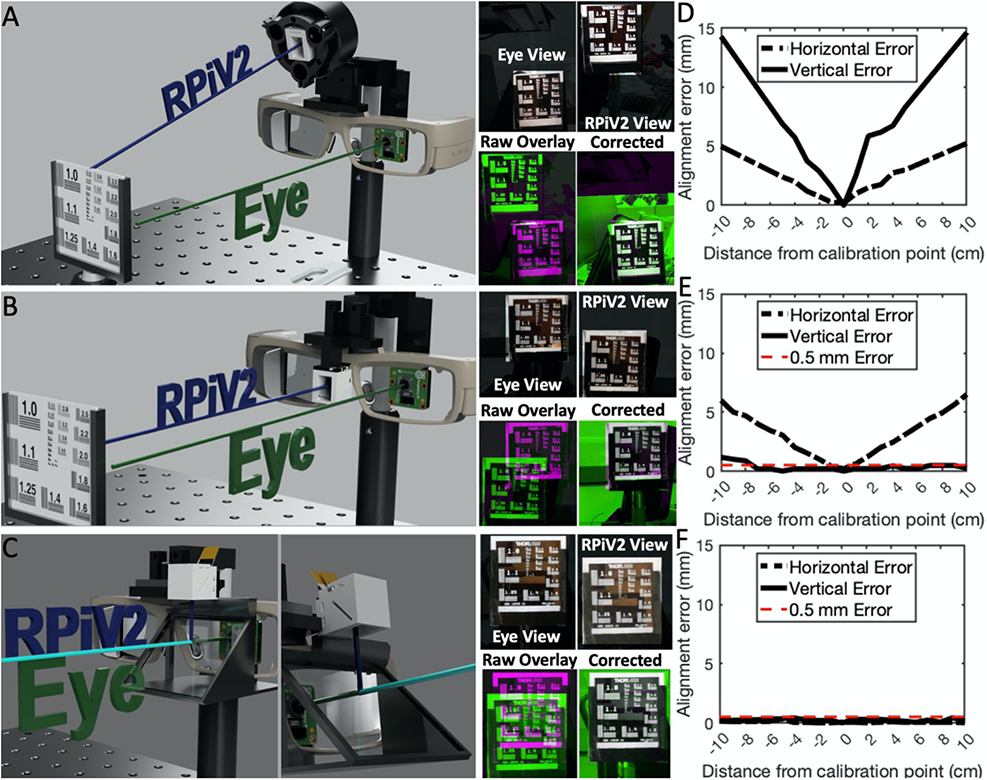
Alignment between FAR-Pi imaging module camera and surgeon’s view can be assured through software-based projective transformation at fixed distance, and can be assured at any distance through optical alignment of camera and eye. A) Experimental setup to test the alignment error of resolution test target images obtained by a surrogate eye camera (’eye view’) and an imaging module (‘RPiV2 View’) positioned within a laser array that is mounted to heads up display (HUD) glasses. Image overlay (‘Raw Overlay’) highlights image misalignment. A 4-point projective transform defined by corresponding fiducial points between images corrects this misalignment and provides an error-free image overlay (‘Corrected’). B) Repeating the analysis of A) with an alternate design, where the imaging module is placed at eye level between the eyes. C) Repeating the analysis of A) with another design, where a glass beamsplitter is placed in front of the HUD glasses oriented 45 degrees relative to the eye light path, and the imaging module is positioned above the beamsplitter in line with the reflected light path. In this arrangement the light paths for imaging module camera and eye surrogate camera are coaxial. D-F) Horizontal and vertical alignment error between the transform-corrected imaging module camera image and eye surrogate camera image as the distance between eye surrogate camera and resolution target varies from 40 cm to 60 cm (±10 cm from the calibration distance). The results in D, E, and F correspond to the experimental setups in A, B, and C respectively.

The ability of the calibration-distance defined projective transform to correct image misalignment at distances beyond the calibration distance was tested next. In the first design with imaging module centered within the illumination module, for every 1 cm deviation in imaging distance from the calibration distance, vertical and horizontal alignment error increased by approximately 1.5 mm and 0.5 mm (maximum 15 mm and 5 mm error when 10 cm away) respectively (Figure 7D). For the second design with imaging module centered at eye level, every 1 cm deviation in imaging distance from the calibration distance resulted in a vertical and horizontal alignment error increase of 0.1 mm and 0.5 mm (maximum 1 and 5 mm error when 10 cm away) respectively (Figure 7E).

In the setting of a surgical procedure where the distance from surgeon to surgical field varies by less than a few centimeters, the distance-specific alignment errors outlined in Figure 7D-E may be tolerable. However, to make the system more robust and support error-free image alignment over a larger working distance, an additional design was developed where the FAR-Pi imaging module and eye are made to be optically coaxial. In this design a 45 degree beamsplitter (50/50 plate beamsplitter #43-359, Edmund Optics) is added in front of the HUD glasses and the FAR-Pi imaging module is placed above the beamsplitter in line with the reflected light path (Figure 7C). Though the FAR-Pi imaging module and surrogate eye camera are designed to be coaxial, images obtained by the FAR-Pi imaging module and surrogate eye cameras were still misaligned (Figure 7C, ‘RPiV2 view’, ‘eye view’). This misalignment is less significant than in the other designs (Figure 7A, 7B), and as was the case with the first two designs, a 4-point projective transform corrects the misalignment between images, allowing for error free image overlay at a calibration distance of 50 cm (Figure 7C, ‘Corrected’). It is important to note that the alignment error in this design likely reflects inaccuracies inherent in the 3D printed components used to secure the components, and this error is therefore independent of imaging distance. Indeed applying the calibration-distance derived transform to FAR-Pi imaging module images obtained at distances from 40 to 60 cm resulted in maximum vertical and horizontal alignment error smaller than 0.5 mm and 0.3 mm respectively (Figure 7F), which is 10 times smaller alignment error than what was observed with the alternative designs.

### Combining all components into a fully wearable system

Because the redesigned FAR-Pi illumination, imaging, and computational modules are compact and light, we sought to combine them with off-the-shelf HUD see-thru glasses (DK52, Lumus, Ness Ziona, Israel) into a single wearable head-mounted system. Using custom 3D printed adapters, all components were mounted securely to a flexible adjustable headstrap sourced from an inexpensive dental headlamp (Figure 8). 3D printed computational module enclosures were designed for both the RPiV4 implementation, where a Raspberry Pi Zero must be included to convert the second RPiV2 camera into a USB-port compatible camera (Figure 8, right panel), and for the RPiCM4 implementation where both RPIV2 cameras could be directly connected to the RPICM4 without the need for a Raspberry PI Zero (Figure S9). The resulting FAR-Pi assembly was evenly balanced, comfortable to wear, and had a total weight of less than 0.67 kg. Further details for integration of all components into the full FAR-Pi system, as well as a parts list with associated costs and sourcing information, are provided in the Supplementary Methods.

**Figure 8.**
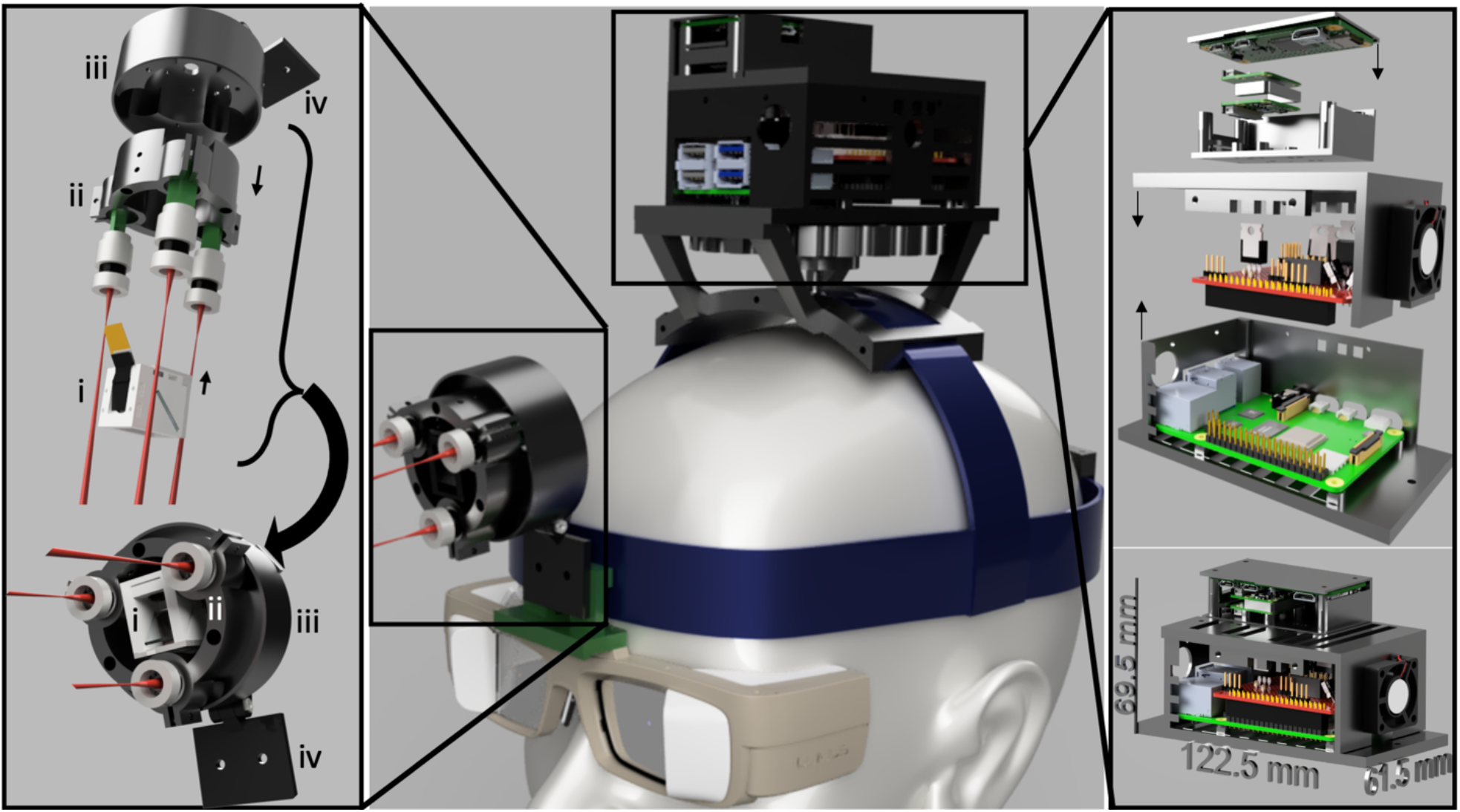
Integrating illumination and imaging modules made from 3D printed and off-the-shelf components with a Raspberry Pi computer on a wearable head mount results in a fully wearable and battery-powered fluorescence-guided surgery (FGS) system. CAD rendering of fully wearable FAR-Pi FGS system with illumination, imaging, computational module, and heads up display glasses mounted to an adjustable surgical headgear. Custom 3D printed adapters are utilized to connect to headgear at forehead and scalp vertex and secure illumination/imaging module and computational module respectively. Left panel shows an expanded view of the imaging and illumination module assembly where the dual vis/NIR enclosure (i) is centered in an enclosure (ii, iii) containing the laser array and an adjustable articulating arm (iv). Right panel provides an expanded view of the computational module components, showing (from top to bottom) Raspberry Pi Zero, 2 RPiV2 PCBs, 3D printed top enclosure with cooling fan, laser diode circuit control HAT, RPiV4, and 3D printed bottom enclosure. Bottom right panel shows a cut-away back-view of the assembled computational module, while center panel shows a front-view.

We found that a 20,100 mAh 5V 4.8A portable USB Anker PowerCore battery power bank was sufficient to power the FAR-Pi system. The current draw from the laser diode HAT with lasers, LEDs, and cooling fan on was <650 mA, the current draw from the Lumus DK52 HUD was <750mA, and the current draw from the RPiV4 with cameras and Bluetooth dongle for user input was <1200mA. This power source provides two separate 2.4A outputs, thus one was used to power the Raspberry Pi SBC, while the other was used to power the laser diode HAT, cooling fan, and Lumus DK52. Given the measured current draw, this system could run for over 7 hours, though if drawing maximum current from the battery the system could operate for over 4 hours. Given that surgeons may use the system intermittently, this time frame is sufficient for many surgical procedures. The portable battery has a size of 16.7 x 6.1 x 2.3 cm and when combined with USB power cables adds a weight of 0.46 kg to the FAR-Pi system. However, the battery adds no head-weight to the FAR-Pi system because it can be comfortably worn on a belt or chest strap.

#### FAR-Pi NIR Fluorescence Sensitivity

The ability of the FAR-Pi system to detect NIR fluorescence with 33 ms exposure (real time imaging) and nM fluorophore concentrations using the FAR-Pi laser diode array was interrogated (Figure 9A). A long-term stable ICG-matching phantom consisting of durable polyurethane with varying concentrations of IR-125 dye was used for fluorescence sensitivity characterization (*28*). Fluorescence signal to background ratio (SBR) was determined for the FAR-Pi system using each of the 4 different excitation filter designs outlined in Figure 3A (Figure 9B). The design with a single 830 nm longpass filter showed the poorest performance, with consistently lower SBR than the alternative filter designs indicating strong background reflected excitation light leaking through the filter. The performance of the single 830 nm filter was particularly poor as IR-125 concentrations decreased below 10 nM – in this regime the decreasing fluorescent signal becomes indistinguishable from the reflected excitation light that leaks through the filter, and the SBR is indistinguishable from 1. The other excitation filter options showed a linear log-log relationship between SBR and IR-125 concentrations from 100 nM to 1 nM followed by a flat plateau at concentrations lower than 1nM. The inexpensive (<$80) double 830 nm filter design showed similar performance to the more expensive 832 nm Edmund BP filter (>$220) and 808 nm Semrock LP filter ($375).

**Figure 9.**
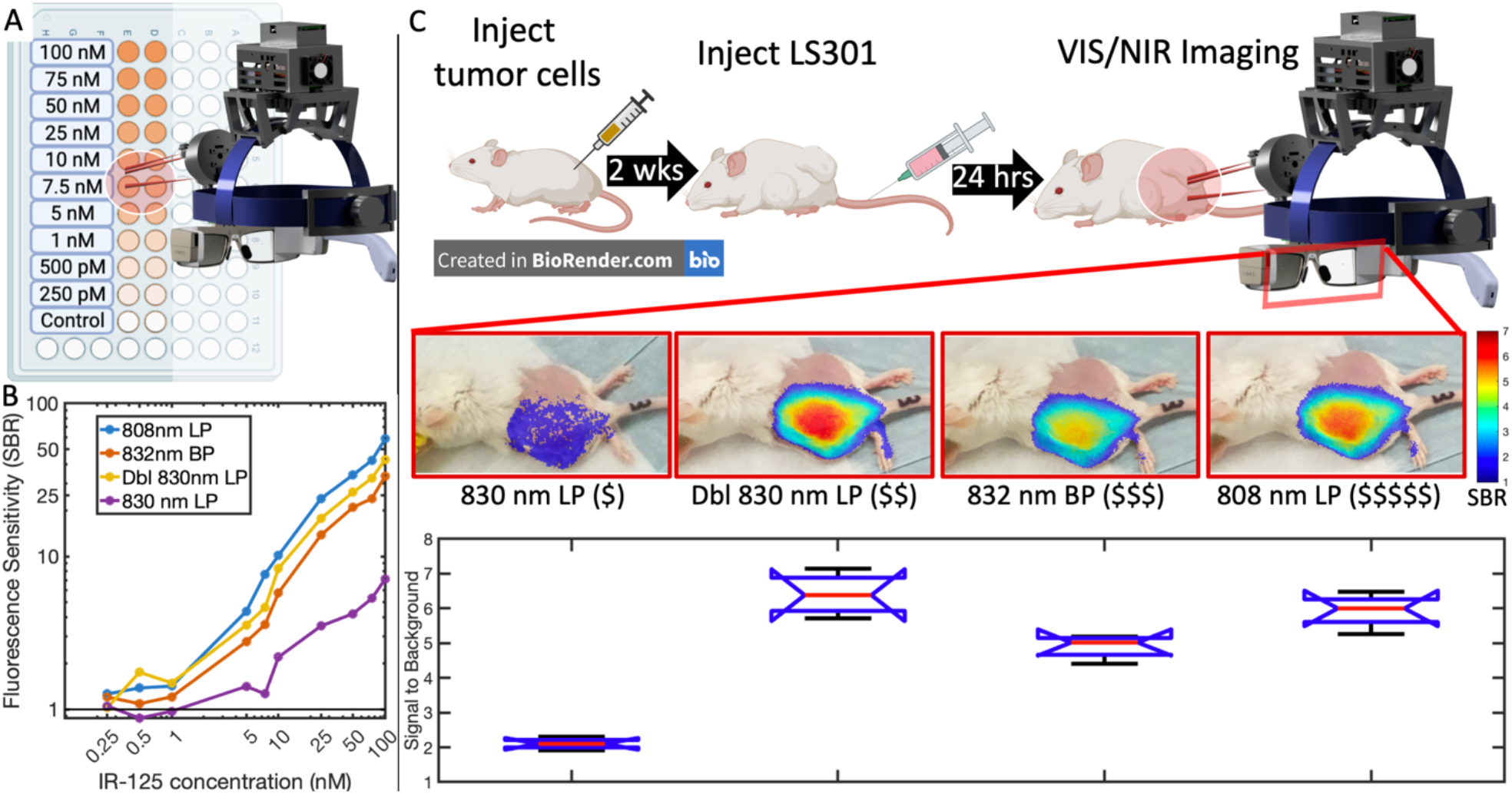
FAR-Pi is sensitive to 1nM fluorophore concentrations and can detect cancerous tissue in an in vivo mouse model. A) Schematic of experimental determination of FAR-Pi fluorescence sensitivity limits using a 96 well plate with IR-125 concentrations ranging from 250 pM to 100 nM. The difference between images obtained with laser on and laser off defined an NIR-background-subtracted image for each well. Fluorescence signal to background ratio (SBR) was calculated for each well by dividing the average background-subtracted pixel intensity of a given well by the average detected background-subtracted pixel intensity in the control well. B) A log-log plot of SBR vs IR-125 concentration for 4 different excitation filter options outlined in Figure 3A: a single 830 nm Newport longpass filter (LP), two 830 nm LP filters in series (Dbl 830 nm LP), one 832 nm Edmund bandpass filter (BP), or one 808 nm Semrock LP filter. C) In vivo validation in a mouse model with left flank subcutaneous breast cancer tumor demonstrating NIR fluorescence from LS301-HSA measured with each of the FAR-Pi imaging module excitation filter options. The $ signs next to each filter name reflect relative expense. In each case co-aligned simultaneous visible and NIR images captured with laser on and off, the NIR signal is processed to generate an NIR-background subtracted image, and then the background subtracted image is divided by an excitation filter-specific background intensity to generate an LS301 SBR heatmap. The LS301 SBR heatmap is shown as a partially transparent overlay on the coaligned visible image using a threshold of SBR>1.5. Lower panel is a box and whisker plot quantifying the peak LS301 SBR value across all mice grouped by the type of excitation filter used during data acquisition. Median is shown in red, top and bottom lines show 75^th^ and 25^th^ percentile respectively. The blue notch indicates a 95% confidence interval around the median.

### In vivo evaluation

The ability of the FAR-Pi system to detect a cancer-targeting fluorophore (LS301) in vivo was tested using a subcutaneous breast cancer mouse model in *n*=4 mice (Figure 9C). Each of the 4 different excitation filter designs described in Figure 3A was evaluated to assess how excitation filter choice impacts fluorescence signal to background. When imaging a white piece of paper, the intensity of excitation light detected in the NIR-subtracted images decreased as the OD of the given filter improved. OD for each filter assembly is shown in Figure 3A. For the 830 nm filter with OD 3-4 near the excitation peak, the maximum pixel intensity of leaked excitation light (out of 1) was 0.53. For the double 830 nm LP assembly, 832 nm BP filter, and 808 nm BP filters the OD around the emission peak was significantly higher and the maximum pixel intensity of leaked excitation light was 0.082, 0.106, and 0.090 respectively. These maximum pixel intensities were used as the ‘background’ in the determination of LS301 SBR for each animal imaged with a given excitation filter assembly. Figure 9C shows a coaligned overlay of the visible camera image with the NIR-camera-derived LS301 SBR, where only pixels with NIR SBR exceeding 1.5 are shown. For each filter the peak LS301 SBR appeared to localize to the center of the tumor.

The bottom panel of Figure 9C demonstrates a box plot of the LS301 peak SBR measured with 4 different excitation filter options across all mice. Data with non-overlapping notches have differing medians (with 95% confidence). For the images acquired with a single 830 nm LP filter, the background laser excitation leak is prominent and as a result the average LS301 peak SBR signal across the 4 mice was significantly lower (2.11±0.16) than the LS301 peak SBR measured with the double 830 nm LP filter (6.40±0.62, *p* = 5.5•10^-8^), the 832 nm BP filter (4.9±0.35, *p* = 6.0•10^-6^), or the 808 nm LP filter (5.93±0.50 *p* = 2.0•10^-7^). The LS301 peak SBR measured with the double 830 nm LP filter was also significantly higher (*p* = 0.0021) than the LS301 peak SBR measured with the 832 nm LP filter but not significantly different (*p* = 0.47) from the LS301 peak SBR measured with the 808 nm LP filter. Finally, the LS301 peak SBR measured with the 808 nm LP was significantly higher than the LS301 peak SBR measured with the 832 nm BP filter (*p* = 0.029).

## Discussion

We previously introduced the GAINS system as an ergonomic real-time FGS system that overcame limitations of bulky, expensive, cart-based fluorescence imaging guidance systems by reducing cost, complexity, and hardware footprint(*17, 18, 29*). Other groups have introduced complementary innovations, including multispectral sensors that obviate the need for separate white light and NIR cameras(*19, 25*), stereoscopic multimodality systems(*21, 22, 27, 30*), and optical methods for ensuring coalignment between the surgeon’s eyes and camera FOV(*31*). Despite the innovations introduced by the GAINS and other more recent systems, FGS HMD systems continue to be wall-powered, complex, expensive, relatively bulky, and replete with significant mismatch between the camera and the surgeon’s view, which makes their adoption in low-resource settings more challenging. To overcome these challenges we leveraged the tools of the “Maker Movement”(*32*) to create the fluorescence imaging augmented reality Raspberry-Pi based goggle (FAR-Pi) system, which to our knowledge is the first battery-powered FGS system where all components (computation module, imaging module, and illumination module, heads-up display) are compact and wearable (Figure 10).

**Figure 10.**
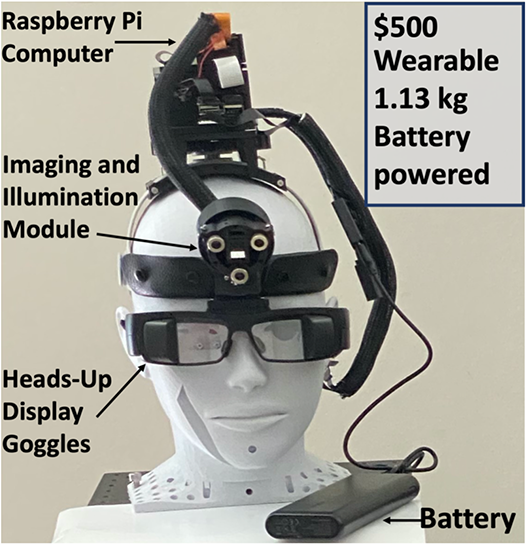
FAR-Pi system demonstrates 10x improvements in cost and weight compared to prior system. All components of the redesigned FAR-Pi system worn on a mannequin head. Apart from the battery pack which can be worn on the hip, the entire FAR-Pi system is integrated into a wearable 0.67 kg head mounted assembly. Excluding the head mounted display, the FAR-Pi system can be built with 3D printed and off-the-shelf components for approximately $500. Unlike previous systems that are orders of magnitude more expensive and bulky, the FAR-Pi does not require wall power, is fully wearable, and does not tether the user to a cart.

Incorporating simple battery-powered low-cost laser diodes in the FAR-Pi system allows the excitation source to be head mounted while also liberating the surgeon from being optically tethered to a bulky or wall-powered light source. The use of flexible MIPI connection cables allows the IMX219 imaging sensor and lens housing to be separated from the larger Raspberry Pi V2 camera module PCB. Separation of the sensor from the PCB allows for close positioning of two imaging sensors (standard and NoIR RPiV2 sensors) and a 45° NIR beamsplitter, thereby achieving dual white light and NIR imaging in a compact (21 mm x 17.625 mm x 16 mm) enclosure. The use of laser diodes instead of broad band LEDs or halogen lamps, as well as the built in IR filter of the standard Raspberry Pi V2 camera, allows the FAR-Pi system to achieve white light imaging free of excitation light contamination without the need for additional shortpass filters. Incorporation of the RPiV4 or RPiCM4 SBC provides a wearable and battery powered Python-based computational engine that efficiently controls laser illumination, dual camera streaming, imaging processing, and image display. Python is a popular open-source software programming language, and building the FAR-Pi system using the Python ecosystem further reduces the barriers to implementing the FAR-Pi system, opening the system for further modification and customization by end-users. The use of a Raspberry Pi Zero running the open-source ‘showmewebcam’ software converts one of the RPiV2 CSI camera into a UVC compliant USB camera, allowing an RPiV4 which has only one CSI camera port to connect to two RPiV2 cameras. The FAR-Pi system is built from inexpensive off-the-shelf components and 3D printed parts and can be constructed without specialized tools or facilities. We also demonstrate that the FAR-Pi system has multiple implementations including different excitation filter options, Raspberry Pi SBC options, laser illumination modules, and approaches to aligning the processed NIR image with the surgeon’s view. Our results demonstrate significant leaked background excitation light leading to limited fluorescence sensitivity and reduced in vivo SBR associated with the single 830 nm LP filter (Figure 9), which may limit the utility of the single 830 nm filter design to only certain use cases where fluorophore concentration is relatively high. However the alternative FAR-Pi implementations overcame this limitation, and interestingly, the cheapest of the alternative filter options 830 nm LP filters) has the strongest in vivo LS301 SBR signal. Finally, the modularity, flexibility, and simplicity of the FAR-Pi system streamlines system maintenance, which is often a challenge for novel technologies in the global health setting(*10*).

The FAR-Pi system is not the first to utilize a Raspberry Pi SBC in an effort to make a more globally accessible FGS system. Li et al developed a battery powered projective imaging FGS system that incorporated a laser diode array, CMOS camera, Raspberry Pi SBC, and a visible light projector to excite and detect near infrared fluorescence from target tissue and then project it with visible light onto the surgical field (*33*). However, while the FAR-Pi system captures real time coaligned visible and NIR images, the Li et al projective imaging system requires exposure times of 330 ms and uses a single camera to detect NIR only. In addition, instead of using inexpensive Raspberry Pi cameras and optical elements like the FAR-Pi system, the Li et al system uses a $400 monochrome USB camera as well as nearly $900 in optical components to ensure appropriate fluorescence detection. In a different application, Kim et al utilized a Raspberry Pi SBC and a RPiV2 NoIR camera to detect parathyroid autofluorescence in real time during surgery, but this system did not capture simultaneous white light images and relied on an expensive high power laser diode source(*23*). To our knowledge, the FAR-Pi system is the first wearable dual visible/NIR FGS system to leverage both Raspberry Pi computers and inexpensive Raspberry Pi cameras for an improved compact form factor.

While the FAR-Pi system has many advantages over existing systems, there are multiple limitations that can be improved upon. The laser diodes have a weak but long NIR tail that contributes to a weak but non-zero background signal which overlaps with the emission peak and thereby decreases SBR. While the effect is negligible at fluorophore concentrations exceeding 1 nM, FAR-Pi system sensitivity to sub-nM fluorophore concentrations could be improved with the additional of excitation clean-up filters (see Supplementary Methods, Section 6). However, the decreased background signal generated by adding clean-up filters must be weighed against both increased costs and decreased signal strength from reduced excitation power. The RPiV2 cameras use fixed focus lenses, and while the depth of focus of the RPiV2 lens ensures appropriate focus for the likely range of distances encountered in practice, implementing automated focus would increase the range of working distances supported by the FAR-Pi system. The illumination module consists of 3 lasers at fixed angle of inclination, with a Gaussian power distribution at the excitation surface. Dynamically adjusting the lasers angle of inclination could ensure a more homogenous power distribution at a given working distance. The FAR-Pi system uses a single NIR camera, and therefore does not capture depth information. A stereoscopic dual visible/NIR imaging system, where there are 2 NIR cameras and at least 1 visible camera, would allow for a more accurate display to the user. Incorporating imaging sensors that implement an RGB-IR Bayer filter with an appropriate notch filter would allow dual visible/NIR imaging with a single camera, but these sensors are typically custom made in research laboratories and not widely implemented in a commercial setting (*19*). Finally, the proposed FAR-Pi system is designed to integrate with a heads-up display device like the Lumus DK52 or Epson Moverio, but these products cost over $1000 and make the total cost of FGS still a challenge for resource-limited settings.

In summary, the FAR-Pi system is an open-hardware inspired, fully-wearable, head-mounted, battery-powered, and easy to build augmented reality FGS solution that is an order of magnitude smaller, lighter and less expensive than existing systems (Figure 10). Incorporation of features that allow laser-camera-surgeon’s view alignment uniquely improves the reliability and capability of the FAR-Pi system, bringing the goal of globally accessible FGS within reach. Further development of the FAR-Pi system are poised to improve human health by making the benefits of FGS available to patients around the world.

## Materials and Methods

### Experimental Design

The objective of the study is to facilitate global access to fluorescence guided surgery by leveraging tools like 3D printing and Raspberry Pi computers and cameras to decrease the clinical footprint, complexity, and cost of existing systems. We consider the key components of existing FGS systems, redesign each component using custom 3D printed and inexpensive off-the-shelf parts and evaluate the functionality of each newly designed component. We integrate the redesigned components together to create the FAR-Pi FGS system, and test the ability of the FAR-Pi system to detect fluorescence from a systemically delivered cancer-targeting fluorophore in vivo in mice.

#### Dual Visible and NIR imaging module

Detection of NIR fluorescence and visible light is a key requirement of FGS systems (*6*). To create a simple, low-cost, and accessible FGS implementation, we first consider the design of a NIR and visible light detection system using inexpensive, customizable, and widely adopted Raspberry Pi SBCs and Raspberry Pi camera modules.

Since 2020, the Raspberry Pi Foundation produced 3 official camera modules, the v1, v2, and HQ cameras that all integrate directly with Raspberry Pi SBCs and support custom image acquisition through python software libraries. These camera modules are inexpensive ($25-30 for v1 and v2, $50 for HQ), available from multiple vendors, and operate seamlessly without additional components. The RPiV2 camera module has higher resolution (8MP vs 5MP) than the v1 camera and unlike the 12 MP HQ camera is available in both visible light and infrared (NoIR) versions. We therefore chose the RPiV2 camera as the basis for the FAR-Pi system (Figure 1B ‘Detection’).

#### Sensor and Lens Specifications

The RPiV2 was first released in 2016 and consists of a Sony 8MP (3280 x 2464 1.12 µm square pixels) IMX219 CMOS sensor that snaps into a camera module circuit board. The circuit board has a standard 15 pin MIPI connector that can be connected directly to a Raspberry Pi SBC camera serial interface (CSI) port. The IMX219 imaging sensor of the RPiV2 camera module is integrated into an 8mm x 8mm x 5mm lens housing. The visible light version of the RPiV2 camera filters out infrared with an IR filter placed between the imaging sensor and lens, while the NoIR version has no IR filter in front of the IMX219 sensor. Though the characteristics of the RPiV2 built-in IR filter have not been released by the Raspberry Pi Foundation, the relative spectral response above 700 nm has been shown to be <1% (*34*). We confirmed this by removing the built-in IR filter from a standard RPiV2 and characterizing the spectral response with a visible-NIR spectrometer halogen light source (Figure S1). Python software libraries provided by the Raspberry Pi Foundation(*35*) provide custom control of image and video acquisition parameters with the option to save 8-bit processed or 10-bit raw Bayer data (*36*).

The IMX219 sensor and housing can be attached to the camera module circuit board directly, or can be attached to the camera module circuit board via IMX219 sensor extension cables (B0186, Arducam, Nanjing China) (Figure 2A). These extension cables are inexpensive ($12.99), readily available from Amazon.com or uctronics.com, and allow the integration of the IMX219 sensor into space-constrained designs by decoupling the sensor from the 25 mm x 25 mm camera module circuit board. The IMX219 sensor housing has a central threaded cylinder with an adjustable lens. The lens has a 3.04 mm focal length, 2.0 aperture, and 62.2° by 48.8° horizontal and vertical field of view (FOV) respectively.

#### Spatial Resolution and Depth of Field Experimental Testing

A backlit Thorlabs NSB 1963A resolution test target (R2L2S1N, Thorlabs Inc, Newton, NJ) was positioned in front of an RPiV2 camera module on a linear rail so that the center of the resolution target aligned with the center of the camera field of view. Images of the resolution test target (Figure 2C) were obtained at distances between 30 cm and 65 cm in 5 cm increments. For any chosen group number, a row vector consisting of the total gray-scale pixel intensity across each column defined the vertical line pairs, and a column vector consisting of the total gray-scale pixel intensity across each row defined the horizontal line pairs (Figure 2D). Local maxima and minima of these 1-dimensional vectors correspond to resolved line pairs, and the highest line pair that demonstrated 5 clear distinct maxima corresponding to all 5 line pairs in a group defined the resolution limit at each distance (Figure 2B).

#### Visible and NIR camera coaligned design

A key component of a FGS system is the co-aligned detection of NIR and visible light, which has typically been achieved in existing systems with the use of optical plate or cube beam splitters (*6, 17*). For the FAR-Pi system, an inexpensive 45° angle of incidence cold IR mirror plate beamsplitter (#62-634 Edmund Optics, Barrington, NJ), which reflects visible light while transmitting NIR light, was used. This cold mirror is made of floated borosilicate, with >95% reflection of visible light and >90% transmission of NIR, has an index of refraction between 1.47 and 1.4650 from 650-900 nm, and has dimensions of 12.5 mm x12.5 mm x 1.1 mm. Less than 5% of the incoming NIR light is reflected off the cold mirror, and though this reflected light is incident on the RPiV2 standard camera dedicated to visible imaging, the built-in IR filter of the RPiV2 standard model camera removes this reflected NIR component (*34*), obviating the need for additional filters. The position of cameras and beamsplitter are determined to ensure equal pathlength for both the visible and NIR imaging arms, which requires taking into account differential pathlength due to refraction through the beamsplitter (Supplementary Methods, Section 4)

While laser diode excitation has a narrow peak at 783 nm, there is a non-negligible long tail that extends to the peak of LS301 emission. An excitation filter that cuts only the peak of laser excitation would allow for maximal detection of NIR-fluorophore emission, but would not block the laser diode tail. Given these trade-offs, imaging modules were constructed and tested with multiple excitation filter options with increasing cost: 1) an optical density (OD) 3-4 12.7mm 830 nm LP Newport colored glass alternative filter ($40, 5CGA-830, Newport, Irvine CA), 2) two stacked 12.7 mm 830 nm LP Newport colored glass alternative filters, 3) a 12.5 mm OD6 Edmund 832 nm BP filter ($240, #84-091, Edmund Optics, Barrington, NJ), and 4) a 12.5 mm OD6 808 nm Semrock EdgeBasic LP filter ($535, BLP01-808R-12.5-D, AVR Optics, Fairport, NY). The bottom panel of Figure 3A provides the optical density of each filter combination as a function of wavelength across the relevant NIR fluorophore excitation and emission spectral range.

A 3D printed enclosure was used to secure the RPiV2 cameras and optical elements in place. The same enclosure design was used for each excitation filter option, though for each case the beamsplitter and optical filter slots were adjusted slightly to accommodate the specific chosen filter (Figure S2, S3).

#### Validating coalignment of captured visible and NIR images

Despite accurate accounting of refraction-associated optical path deviation (see Supplementary methods, Section 4) there are several sources of inaccuracy that may result in mis-aligned visible and NIR images. For example, the IMX219 camera sensors may have small errors in placement relative to the RPiV2 camera housing, and the 3D printed enclosures may have inaccuracies related to 3D printer resolution limits.

To quantify the effect of these potential inaccuracies, the dual camera enclosure was positioned at a distance of 30, 50 and 70 cm from a Thorlabs NSB 1963A resolution test target and images from both the standard and NoIR RPiV2 cameras were captured and exported for further analysis. The resolution test target was backlit with a broadband halogen light source (providing visible and near infrared illumination) to ensure that both the visible camera and the NoIR camera (with one of 4 longpass filter combinations in front) had sufficient illumination for imaging. MATLAB (Mathworks, Natick, MA) scripts were developed to analyze and compare visible and NoIR camera images. Images were converted to grayscale, cropped to include either the vertical or horizontal line pairs of the 1.0 resolution target group, and then reduced into row or column vectors by summing pixel intensities along all rows or columns respectively. The 5 peaks present in these row and column vectors correspond to the x and y pixel coordinates respectively at the centers of the vertical and horizontal line pairs, and the number of pixels between peaks defined a mm to pixel conversion for a given image because adjacent peaks for the 1.0 resolution target group are 1.0 mm apart. Plotting these row and column vectors for both visible camera and NoIR camera images provided a quantitative visualization of the degree of vertical and horizontal misalignment between camera sensors (Figure 3D, Baseline).

#### Software-based correction of sensor disparity

The visible and NoIR images of the resolution test target positioned 50 cm from the dual camera module were loaded into a custom MATLAB script and using the ‘cpselect’ tool 4 corresponding fiducial points were manually selected at the corners of the resolution test target in both images. These points were used as inputs to the ‘fitgeotrans’ function to derive an affine geometric transformation that could be applied to the visible image (using ‘imwarp’) in order to more closely coalign it with the NoIR image (Figure 3D, Corrected). Because the misalignment between cameras is a constant regardless of imaging distance, the affine transform defined by the images at 50 cm applies to images at any distance. While this transform was derived through analysis with MATLAB, the transformation matrix can be applied to the real time RPiV2 visible camera stream using python OpenCV. This transform must be determined once for each dual enclosure that is built, and changes from one build to the next.

### Laser Diode Array Characterization

120 mW 780 nm laser diodes were purchased from Laserlands.net (Besram Technology Inc, Wuhan, China). The spectral profile of the lasers was measured directly with a diffuser and visible-NIR spectrometer (USB2000+VIS-NIR-ES, Ocean Insight, Orlando, FL), adjusting integration time to ensure non-saturated peak signal, subtracting ambient background signal, and then normalizing to peak intensity (Figure 4A).

#### Laser array design and surgical field laser power

A parametric CAD design was created in Fusion 360 (Autodesk, Inc, San Rafael, CA) that positioned any number of laser diodes a distance *d* from the center of a circular array with inclination angle *θ* relative a normal from the laser array center (Figure 4B). In order to have sufficient space for a centrally placed imaging module, *d* was set to a distance of 18.475 mm between laser center and array center. An angle *θ =* 3° 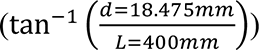 of inclination towards the center of the laser diode array was chosen to support laser intersection at approximately 40 cm. Decreasing the distance *d* brings the lasers closer together and allows for more compact laser illumination module designs (Figure S6).

A laser array consisting of three 780 nm laser diodes designed to intersect at 40 cm and have sufficient space for a central imaging module was constructed (Figure 4D, S4) and aimed at a flat black posterboard target that was positioned at 40 cm, 50 cm, and 60 cm away. The collimating lenses of each laser diode was adjusted to focus the beam waist as close as possible so as to ensure maximum divergence at the surgical field. Maximum power was determined empirically by manually positioning an optical power meter (S121C and PM100D, Thorlabs Inc, Newton, NJ) along the illumination plane until maximum power was obtained. The surface temperature (T) of the posterboard target was measured with a FLIR T650Sc thermal camera (FLIR Systems, Wilsonville, OR) before and 5 seconds after turning on the laser array. The FLIR jpg images were imported into MATLAB (Mathworks, Natick MA) and surface temperature spatial 2D matrix before laser illumination (T(x,y)*_t_*_=0_) and 5 seconds after laser illumination (T(x,y)*_t_*_=5_) were extracted using the FLIR ATLAS MATLAB SDK(*37*). Subtracting T(x,y)*_t_*_=0_ from T(x,y)*_t_*_=5_ yields ΔT(x,y), a 2D matrix containing the spatial profile of temperature increase after laser illumination. Multiplying the ratio ΔT(x,y)/ΔT_MAX_ by measured maximum laser power provided an estimate of the surface laser power spatial distribution (Figure 4C).

### Computational Module

The RPiV4 SBC is light weight (46 g), wearable (88 x 58 x 19.5mm) and inexpensive ($45 for the 2GB RAM model, $55 for the 4GB RAM model) (Figure 1B, Computation top). The RPiV4 has a quad-core 1.5 processor, 2-8 GB of RAM, 2 HDMI display outputs, 4 USB ports, 1 CSI port, and 40 GPIO pins that can be directly integrated with custom external electronics. The RPiV4 hard drive and operating system run on a separate micro SD card. To ensure stability, we used version 10 of Raspbian Buster, a 32 bit legacy Raspberry Pi Foundation operating system based on Debian Linux. This operating system was flashed onto a Sandisk 64 GB class 10 micro SD card ($12.19, ASIN #B08GYBBBBH, Amazon.com) and inserted into the RPiV4.

The single CSI port on the RPiV4 creates a challenge because there are two RPiV2 cameras in the FAR-Pi system that need to be connected to the RPiV4 for simultaneous image capture and processing. There are several options for acquiring synchronized image streams from multiple Raspberry Pi camera modules. One approach is to have multiple Raspberry Pi computers each with CSI-connected Raspberry Pi camera modules that are then synchronized together through an ethernet router (*38*). Another option is using 3^rd^ party hardware add-ons, like the $129.99 Arducam synchronized stereo camera HAT that allows two IMX219 sensor-based cameras to be connected to one CSI port(*39*).

A less bulky, simpler, and more cost effective strategy is to convert the RPiV2 cameras into USB cameras that can be plugged into a RPiV4 USB port (Figure 5 A,B). For example, Arducam offers a $30 UVC complaint USB Camera module (B0196, 8MP 1080P USB Camera Module) that the RPiV2 IMX219 sensor can plug into (item #6 in Figure 5A). This approach relies on smaller 3^rd^ party manufacturer support, and innocuous appearing changes may lead to feature loss. For example, the first version (Rev A) of the Arducam USB Camera Module circuit board supports the use of a camera sensor extension cable (item #4 in Figure 5B), but the firmware on the second version (Rev B) does not. Such incompatibilities can be addressed through direct interaction with the manufacturer, but point out the inherent instability introduced when a design relies on integration of products from disparate hardware providers.

A solution that overcomes this challenge is the open-source “*showmewebcam*” project (*40*), which relies solely on Raspberry Pi Foundation hardware. The *showmewebcam* software converts the RPiV2 camera module into a UVC compliant USB webcam by passing the imaging data through a compact (66 mm x 30.5 mm x 5 mm) and inexpensive Raspberry Pi Zero SBC. Setup consists of flashing the latest “showmewebcam” repository image file onto a >64MB micro SD card, inserting the SD card into the Raspberry Pi Zero, connecting the RPiV2 camera via CSI cable to the Raspberry Pi Zero, and then connecting the Raspberry PI Zero USB port into a RPiV4 (Figure 6A, #7-10). In this configuration the RPiV2 camera stream is detected as a USB webcam, and the total cost is ∼$35-$40. Though this solution has slightly greater physical footprint and is ∼$5-10 more expensive that the Arducam USB camera module, it is arguably a more stable solution because it relies only on Raspberry Pi Foundation hardware and open-source software.

Another approach for dual camera streaming is to leverage the Raspberry Pi Compute module 4 (RPiCM4), which has the same processing speed as the RPiV4 but supports 2 instead of 1 CSI input ports. While the RPiV4 has several input and outputs ports built in, the RPiCM4 has no ports and requires input/output carrier boards to interface with peripherals. It is possible to design a custom carrier board for the RPi CM4(*41*), but there are fortunately several inexpensive premade options that expose both CSI inputs(*42*).

To test the ability of the RPiCM4 to support dual RPiV2 camera streaming, a RPiCM4 with 4GB RAM, 16 GB eMMC and WiFi/Bluetooth chip ($65, CM4104016, Seeed Studio, Shenzhen, China) was plugged into a carrier board ($20, CM4-IO-Base-A, Waveshare, Shenzhen, China) (Figure 5D, item #1), and two RPiV2 camera module PCBs (Figure 5D items #3, #7) were connected to the 2 CSI ports on the carrier board (Figure 5D items 2, 6). RPiV2 standard and NoIR sensors (Figure 5D items #5, #9) were connected to each PCB with Arducam extension cables (Figure 5D items #4, #8). Instead of flashing the operating system to an external microSD as was done for the RPiV4, the operating system (Version 10 of the 32 bit Raspbian Buster) was flashed directly to the onboard eMMC. Setup of the CM4 was more complex than the RPiV4 as it required a 3^rd^ party carrier board and the setup process with the CM4 required more technical expertise to write the operating system to the eMMC. However, the process was well documented in online resources (*43*), and the use of the CM4 allows for native dual camera streaming via CSI port, obviating the need for the Raspberry Pi Zero “*showmewebcam*” workaround. The total cost of implementing the CM4 approach or the RPiV4 approach is equivocal – cost savings realized by removing the need for the RPiV4 micro SD card and Raspberry Pi Zero are counterbalanced by the need for a 3^rd^ party carrier board.

#### Software support for dual camera streaming and display

For the RPiV4-based implementation, custom Python software was developed to process the image stream from both the CSI-connected NoIR RPiV2 camera and the USB-connected standard RPiV2 camera. To read the data from two different cameras synchronously, each camera stream was processed on a separate python thread. The python package ‘picamera’, which serves as a python interface for any Raspberry Pi camera module connected via CSI port (*35*), was used to control image acquisition and camera settings for the NoIR RPiV2 camera. The USB-connected RPiV2 camera could not be directly accessed with the ‘picamera’ package and instead the python OpenCV *VideoCapture* class (*44*) was used to initialize, set parameters for, and read images from the USB-connected RPiV2 camera. To ensure equivalent field of view, both the CSI-connected and USB-connected cameras were set programmatically to have the same resolution of 1920 x 1080. For the NoIR camera this is achieved using sensor mode 1, which supports up to 30 frames per second (fps) without pixel binning. Higher frame rates (40-90fps) are supported by reducing resolution size and allowing 2 x 2 pixel binning.

For the CM4-based implementation, the OpenCV *VideoCapture* class was not necessary as both RPiV2 cameras were connected by CSI input ports and therefore could be detected by the picamera.PiCamera class using the *camera_num* parameter to select either the default camera (*camera_num*=0) or the 2^nd^ connected camera (*camera_num*=1) (*35*). Since the NIR stream does not contain any relevant color information, only a monochrome pixel intensity is necessary. Each frame can be acquired in RGB mode, where pixel intensity must be calculated from R,G,B values separately, or can be captured in ‘YUV’ mode where pixel intensity is in the ‘Y’ channel and color is in the ‘UV’ channel. Using YUV mode is computationally advantageous because it reduces the need for grayscale conversion. Alternatively, both camera streams could be acquired simultaneously by setting the *stereo_mode* parameter of the picamera.PiCamera class to ‘side-by-side’. Each captured frame can then be cropped into left and right halves, each half corresponding to either the NIR or visible camera stream. These halves can be further processed with OpenCV image processing algorithms in real time on a frame-by-frame basis. This ‘side-by-side’ approach has been used previously to generate real time stereoscopic depth mapping with the CM4 (*45*).

In both RPiV4-and CM4-based implementations the Python OpenCV package was also used to perform real-time image processing operations, including image cropping, affine transforms, thresholding, merging, and real time image subtraction. For example, a constant affine transform can be applied in real-time to the USB-connected visible images to correct for the disparity between the visible and NIR-dedicated sensors (see above, *Software-based correction of sensor disparity*).

#### Laser diode control through custom software-controlled electronics

A custom ‘Hardware Attached on Top’ (HAT) circuit was designed to control the laser diode array through RPiV4 GPIO pins. The HAT consists of circuitry constructed on a custom breadboard kit with a 20 x 2 female header (ASIN B08M9QS31F, Amazon.com) designed to be installed directly onto the Raspberry Pi GPIO pins. The circuit (diagramed in Figure 5E) provides power to the 3.3 V laser diode array with a 5V->3.3V voltage regulator (LD1117AV33, STMicroelectronics, Mouser Electronics), 3 LEDs LED (COM-09594, SparkFun Electronics, Niwot, Colorado) that indicate laser on/off status, and a cooling fan (ASIN B08LHBGTL4, Amazon.com) to keep the RPi SBC operating within safe temperature limits. There are two options for powering the circuit – 5V can be supplied from the RPIV4 GPIO pin #2 or #4, which draws power directly from the RPIV4 power supply, or an external 5V power supply connected with a micro USB breakout board #1833, Adafruit Industries, New York, NY) can be used. Both options were implemented in the custom circuit, and switching between options is achieved with a jumper cable. To achieve switching of each individual laser diode and of the entire laser diode array, N-channel MOSFETs (FQP30N06L, OnSemi, DigiKey Electronics, Thief River Falls, MN) were utilized. An additional 10 kOhm pull-down resistor was added between GPIO Pin 25 and ground to ensure that during Raspberry Pi computer startup there was no GPIO control pin switching due to floating voltages.

#### Synchronizing laser excitation with camera frames

For a given region of interest, the difference between image intensity with excitation laser OFF and excitation laser ON generates a fluorescence emission signal where background NIR is removed. Successfully implementing this algorithm on the Raspberry Pi is challenging however, because the RPiV2 camera is a rolling shutter camera that does not have built in support on the camera PCB for detecting when frame exposure starts and stops. However, it is possible to modify the device tree of most Raspberry Pi computers to provide a readout on a selected GPIO pin for when each camera frame is read into memory, and the time at which a frame is read into memory can serve as a delayed surrogate for when camera sensor exposure occurs. We modified the device tree(*46, 47*) in both the RPiV4 and RPiCM4 so that GPIO pin 40 goes HIGH with camera frame read start events and LOW with camera frame read stop events (Figure 6A, camera pulse). Python code was developed to trigger the laser diode master switch (GPIO pin 25) ON at the end of every other frame read event, resulting in laser excitation every 66 ms at 30 fps (Figure 6A, NIR pulse).

To test the ability of this system to synchronize NIR fluorophore excitation and detection, the laser diode array and NoIR camera were connected to the RPICM4 with custom laser diode control HAT and positioned in front of two Eppendorf tubes filled with 1mM indocyanine green (ICG) and water respectively. Laser excitation was pulsed at half the frame rate and synchronized with camera frame capture so that camera frames alternated between having laser excitation ON and OFF. Each frame was captured in ‘YUV’ mode, and only the ‘Y’ channel (the first 3^rd^ of the frame data packet) was used because it contained the grayscale intensity, while the ‘UV’ channel contained irrelevant color information. This reduced the computational burden by a factor of 3 for each frame. We then subtracted consecutive grayscale frames in real time to provide a real time NIR-backgound-subtracted video stream. The start of camera sensor exposure for a given frame may precede the time when the prior frame was read into memory, and therefore when laser excitation is triggered at the end of a frame read event, it is likely starting after frame exposure started. In such a scenario, if the laser remains on for the duration of a full frame it will bleed into the next frame. Thus the duration of laser excitation must be less than the duration of a full frame. We tested setting laser duration to 30%, 50%, 70%, and 90% of frame duration to determine the maximum permissible duration where laser bleed through between frames did not occur.

### FAR-Pi imaging and illumination component integration and image alignment with surgeon’s view

If the FAR-Pi camera is placed in a position that is not coaxial with the surgeon’s eye, there will be misalignment between the camera view and the view from the surgeon’s line of sight. Correcting this misalignment so that the camera view can be superimposed in the HUD with the surgeon’s real-world view can be achieved through software-based image transformation, but the specific image transformation depends on the physical positioning of the imaging module relative to the surgeon’s eye.

The degree of misalignment and durability of software-based realignment, was evaluated the 3 alternative designs shown in Figure 7A, B, and C. For each design a standard RPiV2 camera module was positioned behind the left eye panel of the Lumus DK52 HUD to serve as a surrogate for the surgeon’s real-world view, and then a 1963A Thorlabs resolution target was placed at 1 cm intervals 40-60 cm away from the surrogate eye camera.

We illuminated the resolution target with white light and captured images from the FAR-Pi visible camera and the eye surrogate camera. For each of the 3 designs we extracted a 4-point projective transform between the FAR-Pi image and eye surrogate camera images by using the same MATLAB algorithm outlined in the *Software-based correction of sensor disparity* section. This algorithm can be similarly implemented on the Raspberry Pi SBC in a user-facing GUI by displaying the camera stream in the HUD, having the user select 4 corresponding points between the displayed view and the real-world view, and then using OpenCV (*cv2.getPersepectiveTransform*()) to calculate the projective transform defined by the selected points. The resulting transform can be applied in real time to the imaging module stream to ensure coalignment between the displayed images and the user’s real-world view at the calibration distance. For a given working distance these projective transforms can be set once during an initial calibration step and then can be applied in real time to the FAR-Pi captured image stream to ensure that the images displayed on the see-through HUD glasses are coaligned with the surgeon’s real-world view.

For each design the projective transform defined at 50 cm was applied to resolution test target FAR-Pi imaging module images obtained at working distances ranging from 40 cm to 60 cm, and the vertical and horizontal alignment error between transformed images and surrogate eye camera images was determined by comparing the position of the horizontal and vertical resolution target group 1.0 line pairs (Figure 7D, E, F).

### NIR Fluorescence Sensitivity

To determine NIR sensitivity limits of the RPiV2 NoiR camera, a durable NIR fluorescence phantom was made, consisting of a black 96-well plate (Corning, Corning, NY) with duplicate wells of polyurethane embedded with IR-125 dye in concentrations of 0 (control), 250 pM, 500 pM, 1 nM, 5 nM, 7.5 nM, 10 nM, 25 nM, 50 nM, 75 nM, and 100 nM, as described by Ruiz et al (*28*). A full description of the protocol can be found in the Supplementary Methods, Section 8.

Fluorescence sensitivity of the FAR-Pi system was determined with the IR-125 phantom. Four separate FAR-Pi imaging modules were constructed, each with one of the four excitation filter designs outlined in Figure 3A. These imaging modules as well as a laser illumination module consisting of a 3-laser array were positioned 45 cm above the IR-125 phantom. The imaging and illumination modules remained fixed, while the phantom was placed on customized motorized stage. The illumination module was adjusted so that the first well had an illumination power of 20 mW/cm^2^. The RPiV2 cameras in each imaging module were connected to an RPiV4 computer and images were saved with laser on and off, at multiple exposure times (33 ms, 66 ms, 99ms), with auto white balance on and off, and with ISO set to 400,600, and 800. Images were captured in duplicate for each setting. Then the phantom was moved on the motorized stage so that the adjacent well was positioned at the same location as initial position of the first well. A python script was written to automate laser excitation, image acquisition with different camera settings, and motorized stage positioning. Saved images were analyzed in MATLAB. For each concentration there were 4 images acquired (two wells with same concentration, two images acquired per well). Fluorescence images were converted to grayscale, and mean of pixel intensity within the illuminated well across the 4 acquired images was calculated. The signal to background ratio for each IR-125 dye concentration was defined as the average pixel intensity within each illuminated well region of interest divided by the average pixel intensity in the illuminated control well.

### In vivo studies

All animal studies were performed under an approved protocol by Washington University in St. Louis’s Institutional Animal Care and Use Committee (IACUC). Animals were housed under a 12 h dark-light cycle.

Adult Fox Chase SCID Beige (FCSB) mice (*n* = 4) were subcutaneously implanted with 5•10^5^ 4T1 breast cancer cells on the left dorsal flank. After 2 weeks tumors growth was evidence on the dorsal flank. Each mouse subsequently underwent tail vein injection of 100 µL of 60 µM LS301-HSA solution, which is an NIR cancer-targeting fluorophore developed in our lab (*26*). After 24 hours, all mice were imaged using the FAR-Pi 3-laser illumination module and 4 different FAR-Pi dual VIS/NIR imaging modules, each implementing one of the 4 excitation filter designs presented in Figure 3A. For each imaging module an NIR-to-VIS projective transform was determined to ensure alignment between NIR and visible images (as demonstrated in Figure 3D). The imaging and illumination modules were positioned perpendicular to and at a distance of 45 cm from the imaging plane. For each imaging module FAR-Pi NIR images with ISO set to 800 and auto white balance off of a white piece of paper with and without laser excitation were obtained to determine background NIR and assess the intensity of leaked laser excitation light. The difference between ‘laser-on’ and ‘laser-off’ images for each FAR-Pi imaging module quantified the intensity of excitation light that leaked through the excitation filter assembly associated with a given imaging module. The maximum intensity of leaked excitation light defined the background pixel intensity used to calculate SBR. Thus detected fluorescence has to have intensity above this value in order to have an SBR distinguishable from 1.

Once background leaked excitation light images were obtained, each mouse was imaged using each of the FAR-Pi NIR imaging modules (4 different excitation filter options). For each imaging module, coaligned visible and transcutaneous fluorescent NIR images were obtained with both laser illumination on and off with 1920×1080 resolution and 33 ms exposure. For the NIR camera ISO was set to 800 and auto white balance was off. For each mouse first an NIR-background-subtracted image was calculated by subtracting the gray scale pixel intensity of the laser-off image from the laser-on image. Then the intensity of each pixel was divided by the maximum background pixel intensity. This analysis generated a 2D matrix where the value of element *i,j* corresponds to the LS301 fluorescence SBR at that pixel. This matrix was scaled uniformly to create an LS301 SBR heatmap. The maximum value of LS301 SBR across all pixels defined the peak LS301 SBR for each mouse under each imaging module filter option. In order to exclude signal from non-specific LS301 binding, a mask was applied to this image which excluded pixels with SBR less than 1.5, and then the image was overlaid with partial transparency over the visible FAR-Pi image. Prior to image overlay the NIR-to-VIS projective transform was applied to the NIR-derived LS301 SBR heatmap (Figure 9C middle).

### Statistical Analysis

Descriptive statistics were used. For the in-vivo studies, the peak LS301 SBR values for each mouse were compared between each of the 4 imaging modules with one way analysis of variance (ANOVA), using the *anova1* and *multcompare* functions in MATLAB.

## Acknowledgments

We are grateful for Julie Prior and Brad Manion for assistance with animal experiments in the Molecular Imaging Center. We thank Rui Tang for LS301 synthesis. We thank Xiao Xu for helpful discussions related to illumination module design.

## Funding

National Institutes of Health grant R01EB030987 (SA, CO)

Dermatology Foundation Dermatologist Investigator Research Fellowship (LS)

National Institutes of Health T32EB021955 (LS)

Siteman Cancer Center Small Animal Cancer Imaging Shared Resources:

National Institutes of Health - S10 OD027042

National Cancer institute - P30 CA091842

## Author contributions

Conceptualization: LS, CO, SA

Methodology: LS, CO, KN

Investigation: LS, CO

Visualization: LS

Supervision: LS, CO, SA

Writing—original draft: LS,

Writing—review & editing: LS, CO, SA

## Data and materials availability

All data are available in the main text or the supplementary materials. Data and related code are available upon reasonable request.

## Supplementary Materials

Supplementary text, Figures S1-S13, and Tables S1-S4 are attached in a supplementary material document.

## Supplementary Materials

### Supplementary Text

#### 1. Determination of Raspberry Pi V2 standard camera built in IR filter spectral response

The Raspberry Pi V2 camera consists of an IMX219 CMOS sensor covered by a plastic rectangular housing. An IR filter is glued to the bottom of the housing, and the top of the housing has a threaded channel in which a lens is secured. The housing, which is glued to the edges of the sensor, was carefully removed with the aid of a blade. The lens was then removed by holding the housing and turning the lens counterclockwise with a lens adjustment tool that is packaged with RPiV2 NoIR cameras. With the lens removed, the housing effectively served as a 6mm aperture for the built-in IR filter. A broadband halogen light source with diffuser and visible-NIR spectrometer (USB2000+VIS-NIR-ES, Ocean Insight, Orlando, FL) were positioned on opposite ends of the RPiV2 IR filter. Background-subtracted non-saturated spectra of the halogen lamp with and without the filter were obtained and used to generate optical transmission curves for the built in IR filter (Figure S1).

**Fig. S1.**
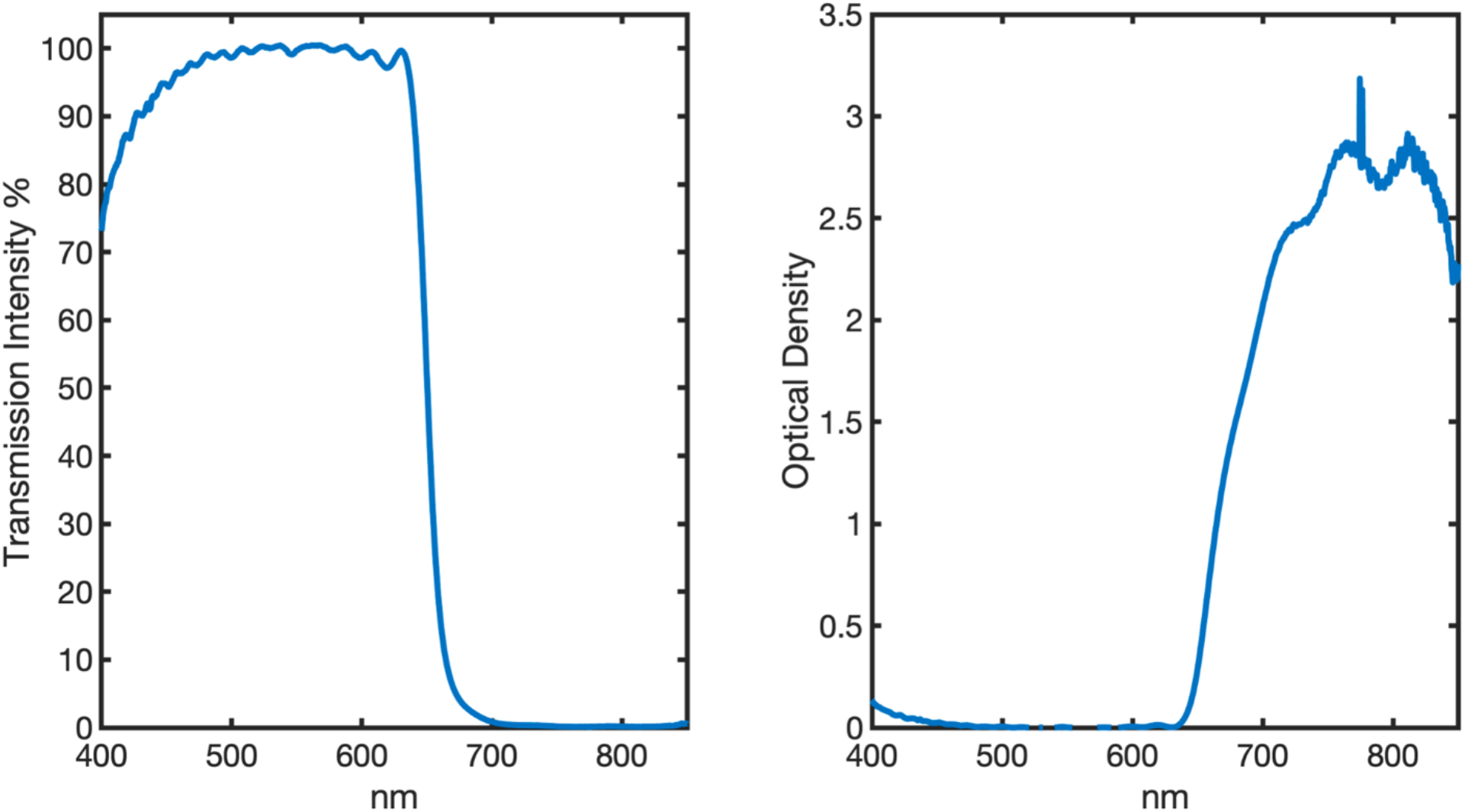
Spectral characteristics of RPiV2 built-in IR filter. Less than 1% of light above 700 nm is transmitted through the filter.

#### 2. RPiV2 camera Spatial Resolution Theoretical Limits

Imaging system resolution is typically expressed in terms of resolvable line-pairs per mm (*lp/mm),* where a line pair consists of 2 contrasting lines (i.e. one light line and one dark line). The theoretical spatial resolution limits at a given imaging distance of the IMX219 sensor can be estimated by first determining the number of pixels per millimeter at a given distance. The pixels per mm along horizontal and vertical direction is given by:

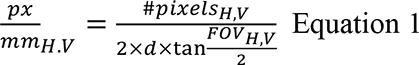

, where *d* is the distance from the camera in millimeters, *#pixels_H,V_* is the number of pixels along horizontal and vertical direction respectively, and *FOV_H,V_* is the horizontal and vertical field of view respectively. Plugging in the sensor dimensions and FOV characteristics of the IMX219 sensor and RPiV2 lens into Eq 1 yields the following expression for both horizontal and vertical pixels per millimeter as a function of distance *d*:

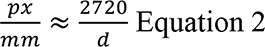

The minimum line-pair thickness is 2 pixels thick, and therefore half of the pixels per millimetre at a given distance defines the maximum *lp/mm* as a function of distance:

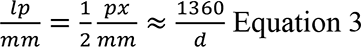

#### 3. RPiV2 Depth of Field Theoretical Limits

The stock position of the RPiV2 lens is focused at a hyperfocal distance of approximately 1 m rendering objects placed at distances of 50 cm to infinity in focus (*48*). The minimum focus distance can be adjusted by manually turning the lens and translating it within its threaded housing to a desired distance from the imaging sensor. Focusing to distances closer than the hyperfocal distance results in a decreased depth of field *Z*’’-*Z*’, which can be estimated from the following equation:

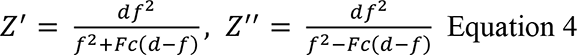

,where *Z*’ is the near-focus distance, *Z*’’ is the far-focus distance, *f* is the focal length, *F* is the aperture, *c* is the diameter of the circle of confusion (largest acceptable blurred sized of a point not in perfect focus), and *d* is the distance where the lens is maximally focused (*49*).

In the FAR-Pi system the camera module can be placed in several positions – in front of the surgeon’s eyes on a forehead mounted articulating arm, between the eyes on the heads up display glasses, or above the glasses with a beamsplitter design. With these different options the distance between camera and surgical field varies from 40 to 50 cm, assuming a working distance of 50 cm between eyes and surgical field. Thus the RPiV2 camera lens must be adjusted from its stock position and be focused closer than the hyperfocal distance, resulting in a reduction in depth of field. Based on Equation 4, focusing the RPiV2 camera to a distance *d* = 40 cm and assuming a circle of confusion *c* equal to the size of three sensor pixels (3•1.12 µm), results in a depth of field of 25 cm, with near-focus and far-focus distances of 31 cm and 56.2 cm respectively.

#### 4. Accounting for Beamsplitter Refraction and Accommodating Alternative Excitation Filters

When using a plate beamsplitter refraction in the transmitted light path results in the optical path being displaced and the optical path length being extended along the transmitted arm (Figure S2). This effect is visible in Figure S2, where the light path from the right that is incident on (*a*) is parallel but displaced a distance (*h*) from the NIR light path (*cd*), and the distance (*ac*) is smaller than the distance the light would have travelled straight through the mirror without refraction (along path *ae*).

Applying Snell’s law, the vertical deviation in optical path assuming an angle of incidence to the beamsplitter of 45° can be expressed by the following equation:

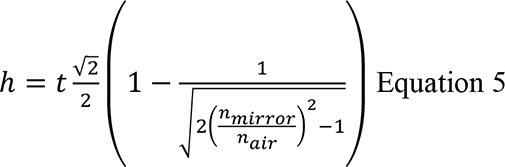

, where *h* is the deviation of the optical path in a direction perpendicular to the incident ray, *t* is the mirror thickness, *n*_mirror_ is the refractive index of the mirror and *n*_air_ is the refractive index of air. For a mirror thickness of *t* = 1.10 mm, *n*_air_ = 1.0003 and *n*_mirror_ varying between 1.4650 and 1.47, the optical path displacement *h* varies from 0.349 to 0.351 mm. The change in optical path length due to refraction *Δl* can be derived and expressed by the following equation:

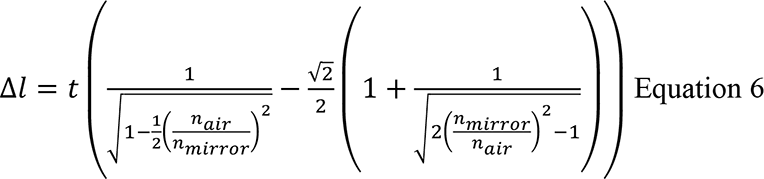

This distance Δ*l* is the difference between distances *ab* and *ae* in Figure 3A, and for a mirror thickness of *t* = 1.10 mm, *n*_air_ = 1.0003 and *n*_mirror_ varying between 1.4650 and 1.47, Δ*l* varies from 0.049 to 0.050 mm.

Applying Snell’s law yields a displacement *h* = 0.35mm and a difference of Δ*l = ab-ae =* 0.05 mm. Thus the use of a plate beamsplitter requires positioning the camera along the transmitted NIR light arm 0.05 mm closer and 0.35 mm higher than would be required if using a cube beamsplitter where there is no beamsplitter-associated deviation of optical path or path length. These deviations in optical path are smaller than the typical errors inherent to 3D printing. To determine suitable placement of cameras sensors and optical elements that met these constraints, a parametric 3D CAD model was developed in AutoDesk Fusion. This model parameterizes the distance *ab* as an adjustable variable, and incorporates Equations 5 and 6 as well as the exact dimensions of the RPiV2 cameras and optical components to ensure that 1) each optical element captures the entire FOV of both cameras and 2) the optical path and pathlength are identical for both cameras. An advantage of this parametric model is that one can change the distance *ab* between camera and mirror to provide sufficient space for optical elements, and the position of both cameras will update programmatically to keep optical path and path length equal.

Setting the distance between the standard RPiV2 camera and cold IR mirror surface *ab* to 5 mm constrains the RPiV2 NoIR camera to have the center *ab*+*h* = 5.35nmm above point *b,* with lens surface at a distance *ae* = *ab-Δl* = 4.95 mm away from point *a*. In this arrangement the transmitted NIR light path is composed of *dc =* 3.755 mm and *ca* = 1.255 mm, which together equals the set 5 mm optical path *ab* between mirror surface and IR-filter containing RPiV2 camera. An excitation light filter is placed in front of the NoIR camera, and therefore there needs to be clearance between the NoIR camera and the beamsplitter edge. To increase the clearance between the edge of the beamsplitter and the NoIR camera, the beamsplitter can be shifted along a 45 degree plane to a limit where the edges of 48.8° degree FOV are incident upon the edges of the beamsplitter (Figure S2B). At this limit with *ab* =5 mm and beamsplitter thickness *t* = 1.10 mm, the clearance *w* is 2.61 mm. The clearance in front and behind the excitation filter needs to be at least 0.1 mm, so the effective space available for the excitation filter is approximately 2.4 mm. This clearance is sufficient for a single 830 nm Newport longpass filter (thickness 1.1 mm), a single 832 nm Edmunds bandpass filter (thickness 2.0 mm), or a single 808 nm Semrock EdgeBasic longpass filter (thickness 2.0 mm). However fitting two 830 nm LP filters with a small air gap is extremely challenging with a clearance of only 2.4 mm, as it leaves only 0.2 mm air gap between filters and leaves no room for the ±0.1 mm thickness tolerance associated with the 830 nm filter. Shifting the cameras away from the beamsplitter increases the clearance for the excitation filter (Figure S2C). To make more space for a double 830 nm filter design, the cameras sensor housings were shifted by 0.4 mm away from the beamsplitter (setting *ab* = 5.4 mm in the parametric design), which increased *w* to 2.88 mm. This width accommodates 0.1 mm clearance between filter and front of the camera surface, 0.1 mm clearance between filter and beam splitter edge, a 0.25 mm air gap between two 1.1 mm thick filters, and leaves 0.23 mm of tolerance for filter thickness.

**Fig. S2.**
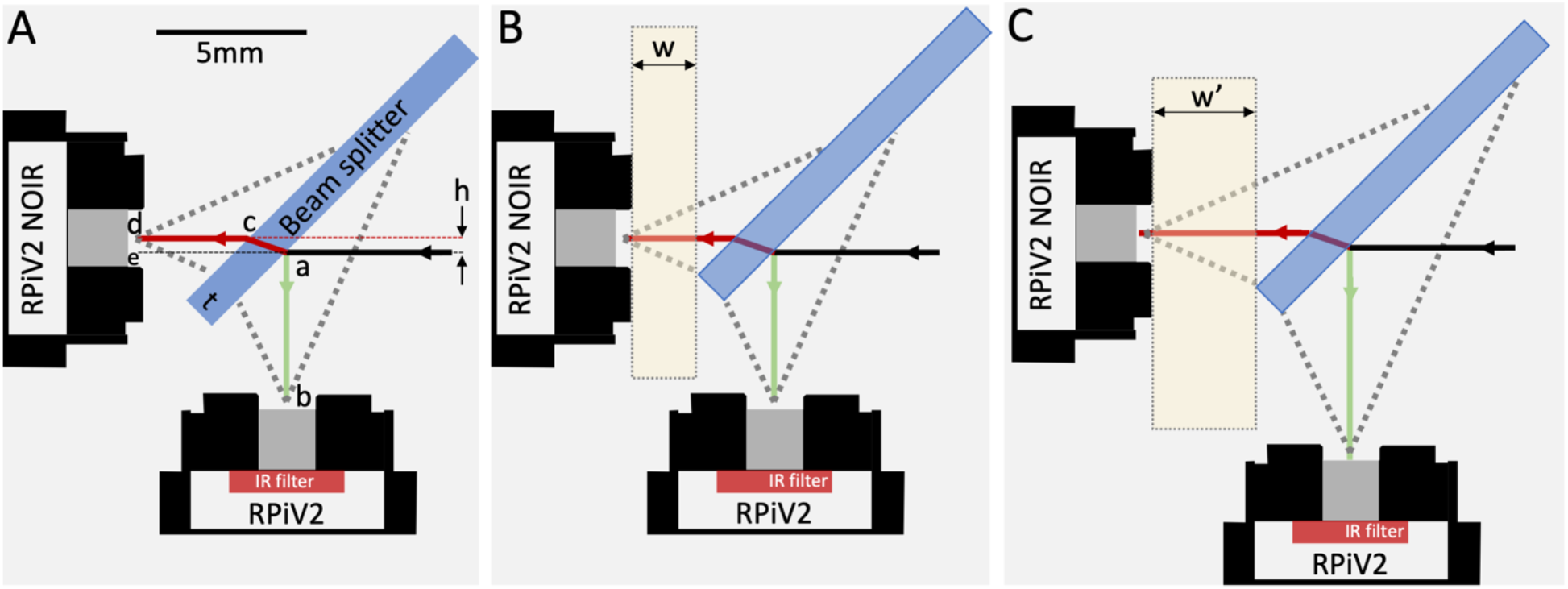
Incoming light (black arrow) is either reflected off the cold IR plate beamsplitter (*ab*) and incident upon a standard RPiV2 camera lens or transmitted and refracted through *(acd*) and incident upon a RPiV2 camera without built-in IR filter. Refraction through the beamsplitter (with thickness *t*) results in vertical deviation *h* of the transmitted light. B) Shifting the beam splitter along the 45 degree plane while ensuring that the lens FOV remains fully incident upon the beam splitter increases the clearance (*w*) between NoIR camera and beamsplitter edge, allowing for thicker long pass filter assemblies to be placed in front of the NoIR camera. C) To increase the clearance to a larger value (*w*’), both cameras can be moved an equal distance away from the beamsplitter. The position of the beamsplitter must also be slightly adjusted to ensure the edges are incident with the edge of the field of view. The cameras in this example were moved by 1.65 mm for demonstrative purposes so that the change in clearance width would be easier to visually perceive.

#### 5. Imaging Module Assembly

A parameterized CAD model was used to create a 3D printed enclosure for the dual RPiV2 camera assembly. The enclosure consists of 3 pieces which were 3D printed on a <$500 CR10v2 printer (Creality3D, Hong Kong) with 1.75 mm PLA (Hatchbox) using a layer height of 0.12 mm and 50% infill. The central enclosure piece has perpendicularly oriented square openings into which the RPIV2 IMX219 sensor housings are press fit, a 45° rectangular slot into which the 12.5 mm square 1.10 mm thick cold IR mirror are pushed into, and vertically oriented slots into which the excitation filters (either single or double) are pushed into (Figure S3A-C).

The imaging module assembly steps are shown in Figure S3 D-E. First the imaging sensor housing from an RPiV2 standard camera and a NoIR RPiV2 camera must be removed from their camera module circuit boards and connected to Arducam MIPI sensor extension cables before being press fit into the 3D printed central enclosure piece. The NoIR camera is positioned on the back while the standard camera is positioned on the bottom. The central enclosure also has eight slots into which M1.4 nuts are pushed into (Figure S3D). Next the cold IR mirror is press fit into the 45 degree central enclosure slot, and the excitation filter is press fit into the vertical central enclosure slot directly in front of the NoIR camera (Figure 3SE). If using the enclosure with 2 slots for 2 excitation filters (Figures S3C), then two filters are slid into two separate slots. In the last step additional 3D printed pieces are added to the back/top of the enclosure and bottom of the enclosure to secure the cameras and excitation filter. Both pieces have 4 M4 holes that align with the 4 M4 holes on the back and bottom of the central enclosure. A total of 8 M1.4 screws are used to secure these two enclosure pieces to the central enclosure.

**Fig. S3.**
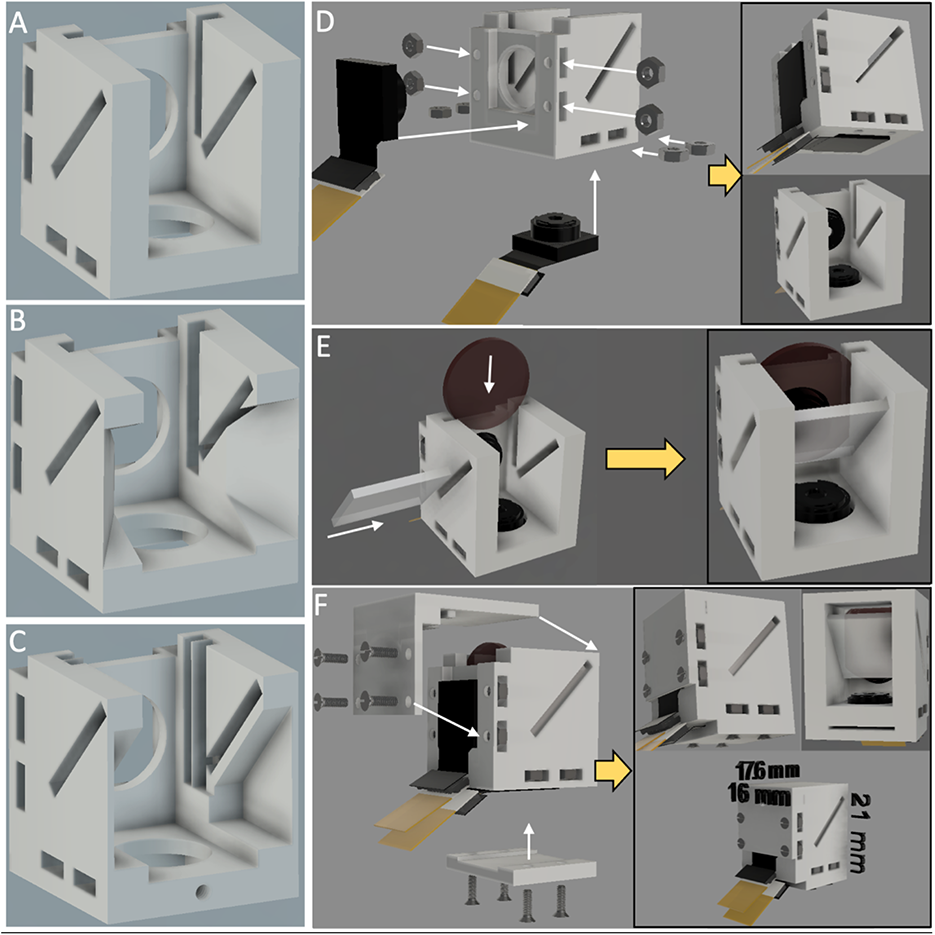
Different designs for imaging module central enclosure supporting alternative excitation filters and step by step assembly all imaging module components. A) Baseline model for central enclosure, supporting a single 1.1 mm thick excitation filter. B) Alternate central enclosure module that supports single 2.0 mm thick excitation filters. In addition some of the internal enclosure wall is thinned out to ensure that camera field of view is not blocked by enclosure wall. C) Alternate central enclosure module supporting two 1.1 mm thick excitation filters with 0.25 mm air gap. D) Assembly step 1, adding RPiV2 camera sensors connected to flexible extension cables and M4 nuts to central enclosure. E) Assembly step 2, adding 45 degree beamsplitter and excitation filter to slots in central enclosure. F) Assembly step 3, securing cameras, excitation filter, and closing central enclosure with 2 outer enclosure pieces and 8 M4 screws.

#### 6. Illumination module Design and Assembly

Several designs were implemented for the laser array, including 1) placing the imaging module in the center of the laser diode array (Figure S4, S5), 2) positioning the lasers as close as possible to each other within the laser diode array (Figure S6), and 3) placing a white light LED in the center of the laser diode array (Figure S7).

##### Assembly – Imaging module centered within laser array

We first describe the illumination module implementation where the imaging module is positioned centrally within the laser array. Figure S4 details the assembly steps. Each 780 laser diode module consists of a 780 nm laser diode epoxied in a 6mm diameter aluminum tube which has interior threads for securing a collimating lens, and exterior threads that secure to a cylindrical aluminum shell designed to protect the laser driver circuit board. To minimize space and weight of the final assembly, the hollow aluminum outer shell was unscrewed and discarded, leaving the laser driver circuitry exposed (Figure S4A).

A 3D printable laser and camera holder was designed to secure the imaging module and 3 laser diodes. The laser holder was generated by the parametric laser array as described above and merged with the imaging module mounting plate. The laser diodes fit in cylindrical holes whose diameter can be clamped down with an M2 screw. The cylindrical holes have an outer diameter of 5.625 mm, which is large enough to not block the laser beam, but smaller than the 6 mm laser outer shell. Thus after removing the collimating lens assembly, the lasers can be pushed into the cylindrical holes until they hit a flat lip, and then clamped in place (Figure S4B). Once this step is complete, the collimating lens assemblies can be reattached, and the imaging module can be secured to the center of the laser assembly using the same M1.4 screws and M1.4 holes used to secure the imaging module bottom outer enclosure in Figure S3F. With the collimating lenses and imaging module secure, the entire assembly is connected with the aid of 3 M2 screws to a bottom outer enclosure that protects the exposed laser driver circuits and provides mounting holes for an articulating arm (Figure S4C). Each laser diode has a black ground wire and red power wire, and each RPiV2 camera sensor has an Arducam sensor extension cable. All of these wires and cables pass through the back of the bottom enclosure and are secured under wire clamps that are mounted to the bottom enclosure (bottom right panel of Figure S4C). The wire clamps ensure that the wires and cables do not get disconnected from the laser driver circuit boards and camera sensors even if the wiring is pulled. To simplify connecting the laser diodes to a power source, the power and ground wires can be incorporated into a 2 pin JST housing or 2 pin female header connectors.

This design also accommodates the addition of individual laser clean-up filters to each of the 3 diodes. The ring assembly to which the 3 laser diodes are clamped has 3 M4 threaded holes equidistant from the laser diodes. An additional 3D printed component that holds 12.5 mm Thorlabs 785 nm laser line premium bandpass filters (FBH05780-10, Thorlabs, Newark, NJ) positioned to fit on the laser diode collimating lens can be secured with these 3 M4 threaded holes (Figure S4D). These additional clean-up filters reduce the intensity of detected NIR background by removing part of the laser’s weak but long NIR tail.

Assembling the imaging and illumination modules can be achieved with 3D printed parts and simple tools like screwdrivers, tweezers and wire crimpers in less than 1 hour. All components were arranged to be printed together and occupied an area of approximately 11cm x 11cm, which fits on numerous inexpensive 3D printers like the Creality Ender 3 (<$200) or CR10v2 (<$500). Printing all components at once on a Creality CR10v2 printer with layer height of 0.12 mm and 50% infill took 10 hours and required 45g of PLA filament (Figure S5A). All components (3D printed parts, RPiV2 NoIR standard cameras, Arducam camera sensor extension cables, cold IR mirror, 830 nm longpass filter, and fastening hardware) were organized on an 8.5”x11” paper template to help organize step by step assembly (Figure S5B).

##### Assembly – Illumination module and imaging module decoupled

Figure S6 summarizes the assembly steps for the illumination module that decouples the illumination module from the imaging module and places laser diodes in close proximity. This close proximity allows for the option of reducing the excitation source long NIR tail with only a single optical filter, which is a cost effective approach to improving system fluorescence detection sensitivity.

All of the components were modeled in Autodesk Fusion 360 and 3D printed prior to assembly. To construct the laser array, first each laser diode module is deconstructed as shown in Figure S4B, and the hollow outer shell is discarded. Then the 3 laser diode modules are press fit into a 3D printer laser diode holder generated from the parametric laser array as described above. (Figure S6A). This holder is designed to place the laser diode modules at 3 degree inclination essentially in contact with each other. As in Figure S4C, the holes that the laser diode module press fit into have a lip that limits the laser diode modules from being pushed through fully through the holes. After press fitting the laser diode modules, the collimating lens assemblies are screwed back into the laser diode module external shells. Next 3D printed outer enclosures are added to the bottom and top of the laser array holder (Figure S6B). The outer enclosure on the back has 3 M3 screw holes that align with 3 holes on the laser array holder. This outer enclosure protects the laser diode module electronics, and also has an opening at the bottom for the laser diode wires. There is a triangular wire clamp that attaches to the bottom of the outer enclosure with 6 M2 screws which secures the 6 laser wires so that pulling on the wires does not put strain on the laser circuit board connections. An additional 3D printed enclosure attaches to the front of the laser holder array, this enclosure has 3 threaded M3 holes that aligns with the 3 M3 screw holes in the bottom outer enclosure and laser holder array. The front of this enclosure has a space that fits a 25 mm optical filter. In the final assembly step, a 25 mm Edmund 785 nm OD 4.0 10 nm wide band pass excitation clean-up filter (#86-739, Edmund Optics, Barrington, NJ) is positioned in the front of the outer enclosure, and held in place with a 3D printed component with hinged cover that is secured to the outer enclosure with 3 M3 screws (Figure S6C). The fully assembled illumination module fits within a cylinder with 35 mm diameter and 55 mm height.

##### Assembly – Illumination module coupled with central white light LED

While the laser diodes produce some visible light (due to the long tail on both sides of the laser diode peak), the laser spot is faint compared to room light. Visualizing the location of the laser spot is important for the FGS system operator to ensure that they are interrogating the appropriate area of interest and also to ensure safe operation of the system with respect to other people in the vicinity of the FGS system. To solve this problem the parametric laser array design was modified by increasing the distance between lasers to allow a white light LED to be placed in the middle of the laser diode array (Figure S7). White light LEDs have weak but long NIR tails that are stronger than the long NIR tail of the excitation laser source, and therefore an IR filter is placed in front of the white light LED filter out any NIR light emanating from the white light source that may increase the detected background signal.

The assembly steps for this design are presented in Figure S7. Similar to the other illumination module designs, all components were modeled in Autodesk Fusion 360 and 3D printed prior to assembly. Similar to the first steps in Figure S5 and S7, first each laser diode module is deconstructed as shown in Figure S4B, and the hollow outer shell is discarded. Then the 3 laser diode modules, with collimating lens assembly removed, are pushed into the laser diode holder holes and clamped with M2 screws (Figure S7A). The collimating lens are screwed back in to the laser diode modules once the modules are secured in the laser holder assembly. This laser holder has a space in the center for a rechargeable battery-powered white light LED ($32.99, ASIN B07WSQ3N6G, Amazon.com), which consists of a bright 5W white LED placed inside a cylindrical tube with a lens at the end. The white light LED is positioned in the center and flush with the laser holder array and secured with M3 set screws. Next, similar to the steps shown in Figure S4C, an outer enclosure is attached with 3 M2 screws to protect the laser diode circuitry (Figure S7B). The back side of this outer enclosure has wire clamps that secure the laser power and ground wires to protect them from being pulled out of the laser diode PCBs (Figure S7B right panel). Finally, similar to the steps in Figure S4D, a 3D printed component is attached with 3 M4 screws to the laser diode array ring which secures three 12.5 mm 785 nm Thorlabs premium bandpass filters to the laser collimating lens assemblies (Figure S7C). In addition, this 3D printed component secures an IR-cut filter (12.5 mm Square IR Cut-Off Filter, #55-235 Edmund Optics) in front of the white light LED. The fully assembled illumination module fits within a cylinder with 54 mm diameter and 60 mm height.

**Fig. S4.**
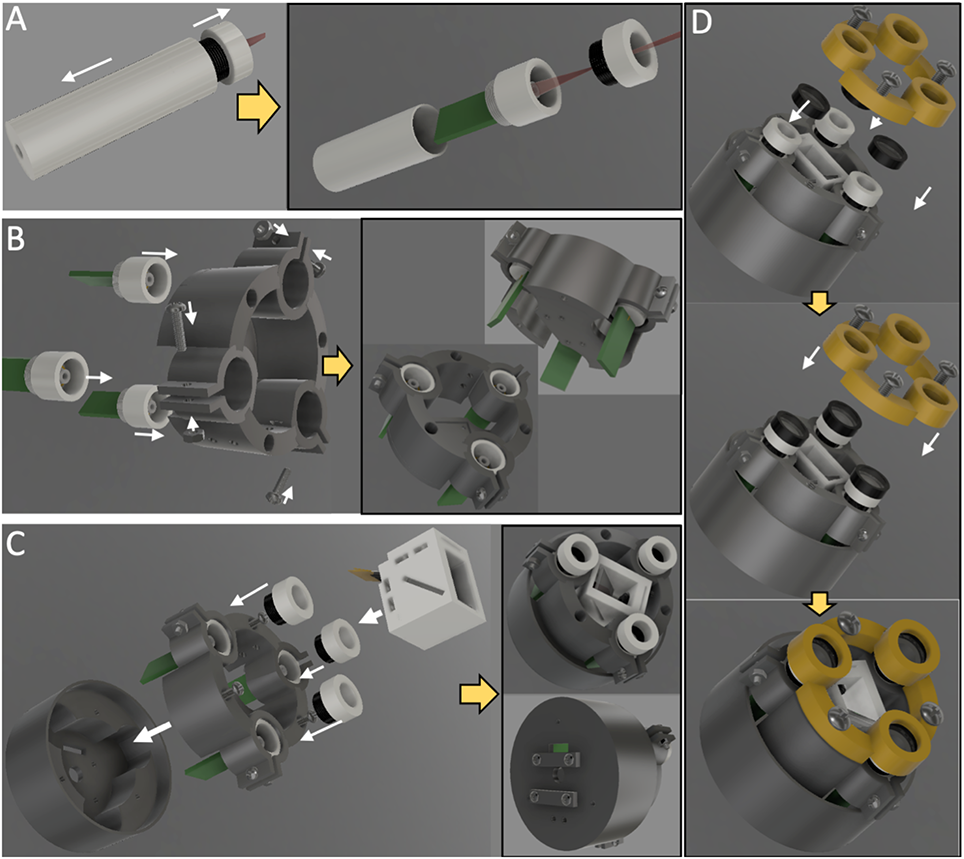
Figure S4. Illumination module with central imaging module assembly and step by step build. A) A single laser diode is first disassembled, the aluminum shell is discarded, exposing the laser diode control circuitry, and the collimating lens assembly in the front is set aside until a later step of assembly. B) Laser diodes are press fit into a custom 3D printed laser array holder. M2 screws and nuts are used to secure each laser diode after it is inserted into the laser array holder. The laser array holder has a space in the middle with mounting holes that line up with the imaging module. C) The imaging module is inserted into the middle of the laser array holder and secured with the same M1.4 screws used to secure the bottom outer enclosure of the imaging module (as in Figure S3F). The collimating lens assemblies are screwed back into each laser diode assembly. The entire assembly is attached to an outer 3D printed laser array enclosure with 3 M2 screws. This outer enclosure protects the laser diode circuitry and also has threaded holes for wire clamps and mounting brackets on the back (lower right panel). D) This design also supports addition of three 12.5 mm 785 nm premium bandpass filters that are fitted on each laser diode and secured with a 3D printed component that attaches to the inner enclosure with 3 M4 screws.

**Fig. S5.**
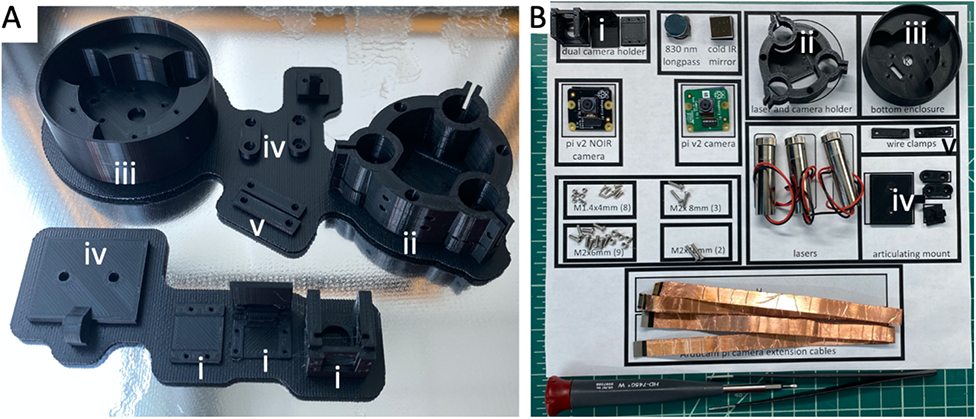
Imaging and illumination module 3D printed components can be printed at once and incorporated with simple tools and off the shelf components into the full assembly. A) 3D printed components printed together, including the imaging module enclosures (i), the laser array and imaging module holder (ii), the laser array outer enclosure (iii), the articulating arm that mounts to the laser outer enclosure (iv), and the wire clamps that attach to the back of the laser array outer enclosure (v). B) All the components necessary to build the combined imaging/illumination module are organized on an 8”x11” template to simplify the build process.

**Fig. S6.**
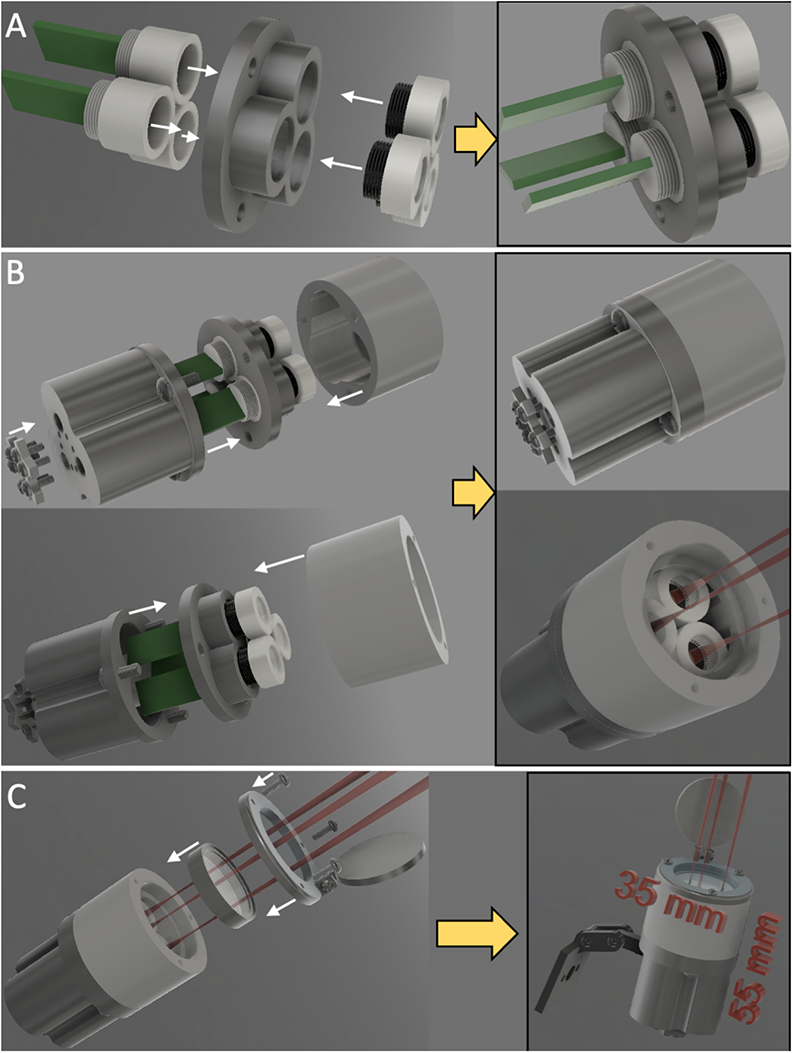
Deconstructed laser diode modules are first press fit into a 3D printed laser holder array, and then the collimating lens assembly are reattached to each laser diode module. B) 3D printed outer enclosure with wire clamp is attached to the bottom of the laser holder array, while a 3D printed outer enclosure and optical filter mount are attached to the front of the laser holder array. Two different views are shown (top and bottom panel) C) A 25 mm shortpass filter is placed into the optical filter mount and secured with using M2 screws with a 3D printed hinged cover plate. An articulating mount can be incorporated with the entire assembly to assist with mounting to surgical headgear (right panel).

**Fig. S7.**
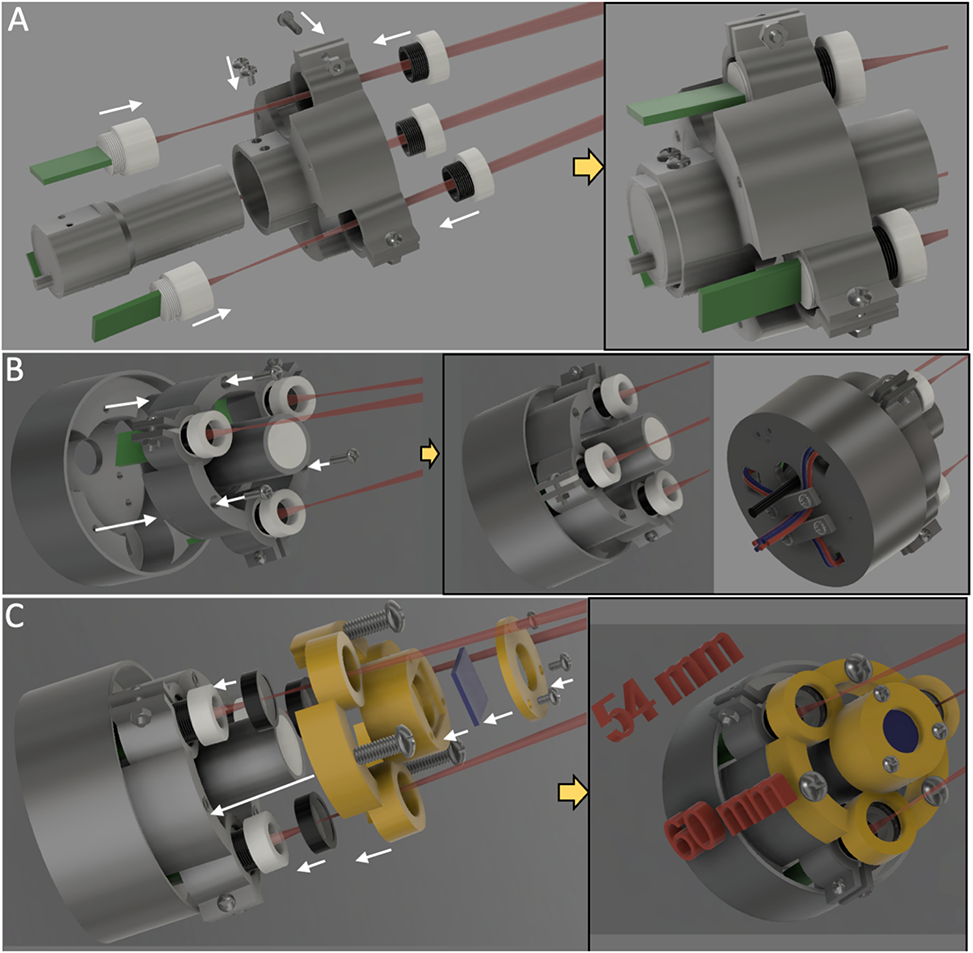
Assembly steps for an illumination module with a white light LED centrally within a laser diode array. A) 3 laser diode modules are positioned and clamped into a 3D printed laser holder array. A white light LED is secured to the center of the laser holder array with M3 set screws. B) The entire assembly is attached to an outer 3D printed laser array enclosure with 3 M2 screws. This outer enclosure protects the laser diode circuitry and also has threaded holes for wire clamps and mounting brackets on the back (right panel) C) This design also supports addition of three 12.5 mm 785 nm premium bandpass filters that are fitted on each laser diode as well as an IR cut-off filter fitted on the white light LED. These optical filters are secured with a 3D printed component that attaches to the inner enclosure with 3 M4 screws.

#### 7. Integration of all FAR-Pi Components

##### Computational module

The computational module can be built either with a Raspberry Pi V4 single board computer (RPiV4) or a Raspberry Pi CM4 single board computer (RPiCM4). In both cases the Raspberry Pi single board computer is combined with a laser diode control circuit, a cooling fan, and camera related hardware. All of these components are housed within a custom enclosure. Figure S8 details the assembly steps needed for building the computational module based on the RPiV4.

First the RPiV4 is connected with GPIO pins to the custom laser diode control ‘Hardware Attached on Top’ (HAT) circuit and the combined RPiV4 and HAT are secured into the bottom of a custom 3D printed computer enclosure. This enclosure is designed to provide access to the RPiV4 input and output ports (Figure S8A). Then the cooling fan and enclosure for the Raspberry Pi Zero and 2 RPiV2 camera module PCBs are attached to the top of the RPiV4 SBC enclosure, which is then secured to the bottom of the enclosure built in the prior step (Figure 8B). Note that the RPiV2 camera module PCBs are attached with a spacer between them, one PCB faces down while the other faces up. The CSI output from the bottom RPiV2 camera PCB connects to the CSI input port of the RPiV4. The enclosure that houses the RPiV2 camera PCBs also has threaded holes for attaching wire clamps. The wire clamps secure the flexible MIPI cables that connect the RPiV2 camera sensors to the RPiV2 camera PCBs. Finally a Raspberry Pi Zero is secured to the top of the enclosure and an outer cover is attached (Figure S8C). This outer cover protects the Raspberry Pi Zero while keeping the CSI input port and USB output port of the Raspberry Pi Zero accessible. The Raspberry Pi Zero CSI input port connects to the top RPiV2 camera PCB CSI output, and a USB cable connects from the Raspberry Pi Zero to one of the RPiV4 USB input ports.

Figure S9 details the assembly steps for building the RPiCM4-based computational module. Unlike the RPiV4, the RPiCM4 has no accessible ports, and must be plugged into a custom carrier board. A $20 CM4-IO-Base-A carrier board (Waveshare, Shenzhen, China) with two CSI outputs exposed was utilized. The carrier board has the same GPIO pins as a standard RPiV4, so the laser diode control HAT can be installed via GPIO pins as well. In the first step of assembly the RPiCM4 is connected to a carrier board, the HAT is connected to the GPIO pins, and the combined components are secured into a custom 3D printed enclosure that exposes the ports of the carrier board. A cooling fan is secured to the enclosure as well (Figure S9A). Since the CM4 can have two CSI camera connected directly there is no need for a Raspberry Pi Zero to be included in the final assembly. As a result, in the final step only 2 RPiV2 camera PCBs (along with wire clamps) are secured to the top of the computational module enclosure (Figure S9B). Both of CSI outputs of the PCBs are fed directly into the RPiCM4.

##### Surgical headband modification

An existing off-the-shelf surgical headband and white light LED ($56.99 B0823QZSLL, Amazon.com) with adjustable headband and overhead strap was used as a template onto which FAR-Pi components could be incorporated. The surgical headband has a plastic component positioned on the mid scalp with a central knob that can be turned to move the vertically oriented flexible straps closer or further apart, thereby adjusting the tightness of the overhead straps. To support mounting FAR-Pi components, we disassembled the overhead tightening components and replaced the central channel that the flexible straps move through with a 3D printed channel with the same dimensions but with threaded mounting holes on the side (Figure S10).

##### Mounting computational module to surgical headband

A custom 3D-printed elevated stage with threaded mounting holes was designed to provide a flat mounting surface for the RPiV4 or RPiCM4 above the adjustment knob. With the computational module mounted over the tension knob, it becomes ergonomically challenging to adjust the knob because the module enclosure is in the way. An adjustment gear with edges extending beyond the RPiV4 and RPiCM4 enclosure bases was designed to fit over the existing knob to allow for easy adjustment of strap tension. Assembly steps are shown in Figure S11A,B for mounting the RPiV4-based computational module and in Figure S11C,D for mounting the RPiCM4-based computation module. In both cases, first an oversized adjustment gear press fit to the existing tension knob, and then the legs of the elevated stages are attached to the custom 3D printed channel installed in Figure S10. With the stage secured, the RpiV4 and RPiCM4 enclosure bases are secured to the elevated stages with M3 screws (Figure S11 B, D).

##### Mounting heads up display and illumination module surgical headband

We chose the Lumus DK52 as a example see-thru heads up display to use in implementing the FAR-Pi. A custom 3D printed component (Figure S12A) was designed in Autodesk Fusion to interface with the M2.5 mounting holes along the top of the Lumus DK52 and the 3 mm mounting holes on the front of the headband. The FAR-Pi illumination module articulating arm (‘iv’, Figure S5) was designed to have the same 3 mm mounting holes as the headband, which allowed both the DK52 heads up display and the illumination and imaging module to be secured together with the front of the headband (Figure S12B). With the articulating arm mounted to the heads up display and surgical headband, the illumination module can be easily incorporated. Figure S12C demonstrates the illumination module with central imaging module (detailed in Figure S4) attached to the articulating arm, while Figure S12D and S12E show the illumination modules from Figures S6 and S7 respectively attached to the headgear-mounted articulating arm.

##### Mounting the imaging module

Figure S12C demonstrates an implementation where the imaging module is positioned centrally within the illumination module. In this scenario no additional build steps are required. However, the imaging module can be placed in different positions in the FAR-Pi system, and adjusting the 3D printed adaptor that secures the HUD glasses to the surgical headband supports these alternative imaging module positions.

In Figure S13A the 3D printed adaptor between HUD glasses and surgical headband is modified to support positioning the imaging module at eye level at the center of the HUD glasses. This modified adaptor starts with the design in Figure S12A, but adds a bracket in the front with 4 mounting holes that align with the 4 M1.4 holes on the back of imaging module enclosure. The imaging enclosure is small enough to fit between the eyes without obstructing vision in either eye. In Figure S13B the 3D printed adaptor between HUD glasses and surgical headband is modified to support positioning the imaging module with a beamsplitter, so as to create a shared optical path between the surgeon’s eyes and FAR-Pi cameras. This modified adaptor starts with the design in Figure S12A, and adds 1) a vertical mount with 4 M1.4 holes that align with the back of the imaging module enclosure, and 2) a 45° shelf where a 50 mm square plate beamsplitter secured in front of the HUD glasses. In both of the assemblies outlined in Figure S13 the articulating arm that is used to mount the illumination modules in Figure S12 is also utilized.

**Fig. S8.**
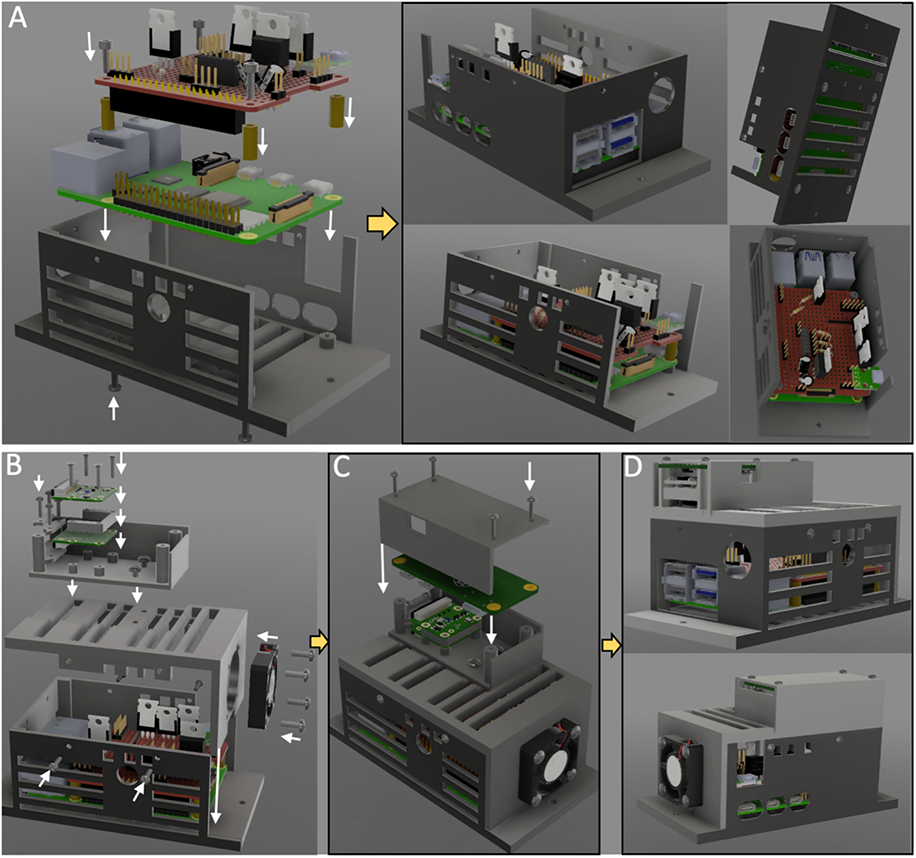
Assembly of Raspberry Pi V4 (RPiV4)-based computational module with enclosure. A) RPiV4 single board computer (SBC) and custom laser diode array controller circuit (red breadboard) are secured with M2.5 screws to a 3D printed enclosure. The 3D printed enclosure has holes exposing the ports on the RPiV4 and controller circuit. B) Wire clamps as well as two RPiV2 camera control boards separated by a 3D printed spacer are secured to a rectangular 3D printed enclosure with M2 screws, and this assembly is secured with M3 screws to a 3D printed component that serves as the top of the computational module enclosure. The bottom RPiV2 camera PCB is connected to the CSI port of the RPiV4 SBC. A cooling fan is secured to the top of the computational module enclosure, and then the top and bottom pieces of computational module enclosure are secured together with M3 screws. C) Raspberry Pi Zero is secured above the RPiV2 control boards with another 3D printed component that serves as the top of an enclosure protecting the Raspberry Pi Zero. The Raspberry Pi Zero CSI port connects to the top RPiV2 PCB. D) The final assembled computational module, with RPiV4 computer handling the RPiV2 NoIR camera stream directly, and Raspberry Pi Zero converting the RPiV2 standard camera stream to a USB camera before passing the camera stream via USB port to the RPiV4.

**Fig. S9.**
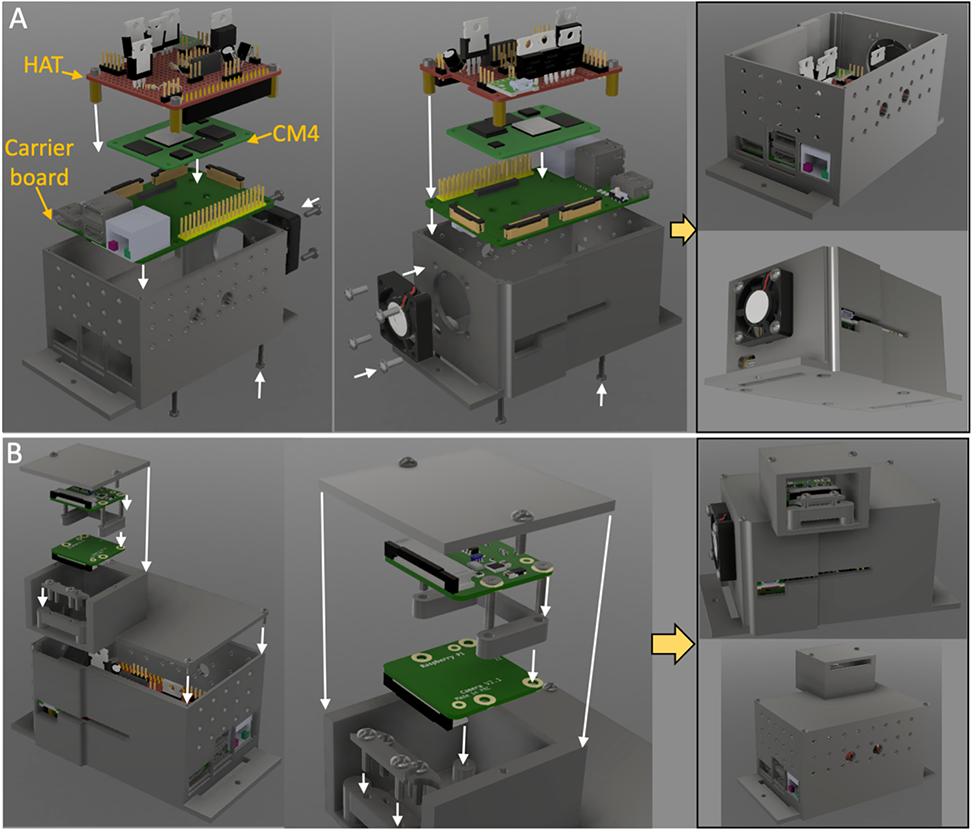
Assembly of Raspberry Pi Compute Module 4 (RPiCM4)-based computational module with enclosure. A) The CM4 is attached to a specialized carrier board before the laser control HAT is attached. All these pieces, as well as a cooling fan, are secured to a custom 3D printed enclosure. B) Raspberry Pi V2 camera PCBs are attached to the top of the computational module enclosure, and the CSI outputs of from the camera PCBs are exposed so that they can feed directly into the two RPiCM4 CSI inputs.

**Fig. S10.**
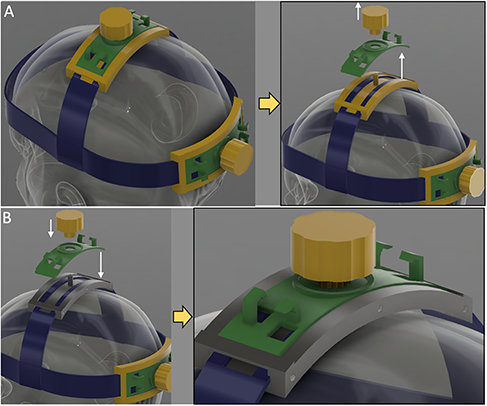
Modifying and off-the-shelf headmount to support mounting additional components to the headmount. A) An off-the-shelf headmount has rack and pinion gear assembly at the scalp vertex that tightens the band that extends from temple to temple over the head. Removing the gear (yellow) and outer cover (green) allows one to remove the channel (yellow) through which the straps (blue) travel. B) The channel (yellow) is replaced with a custom designed 3D printed channel (gray) that has threaded mounting holes on the side. The outer cover (green) and gear (yellow) are then placed back into position.

**Fig. S11.**
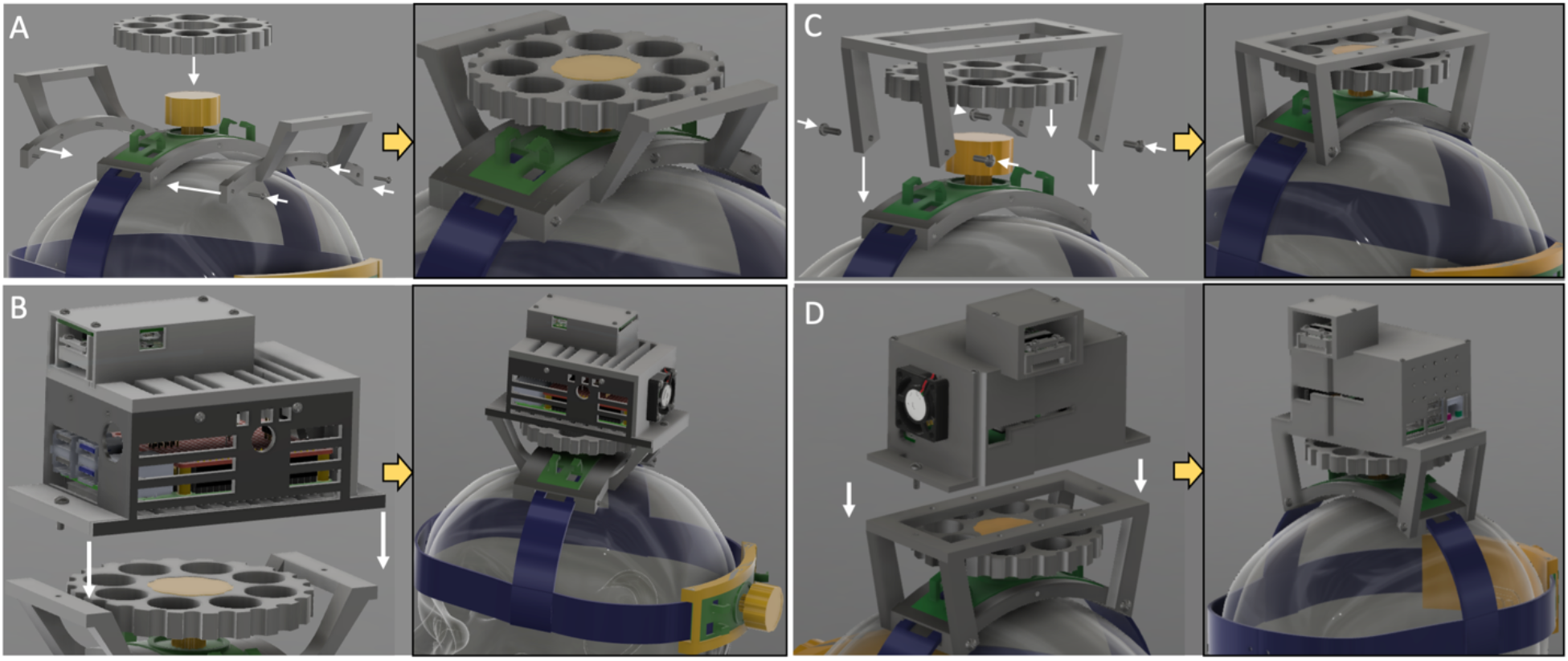
An elevated stage provides a mounting surface for the FAR-Pi computational module enclosure. A) Elevated stage assembly for the RPiV4-based FAR-Pi computational module. B) RPiV4-based FAR-Pi computational module enclosure is secured via M3 threaded holes on the elevated stage. The original tension knob (yellow) is inaccessible with the computation module enclosure installed, but the gear that was installed in step A) makes tension adjustment possible. C-D) Steps A) and B) for the RPiCM4-based FAR-Pi computational module.

**Fig. S12.**
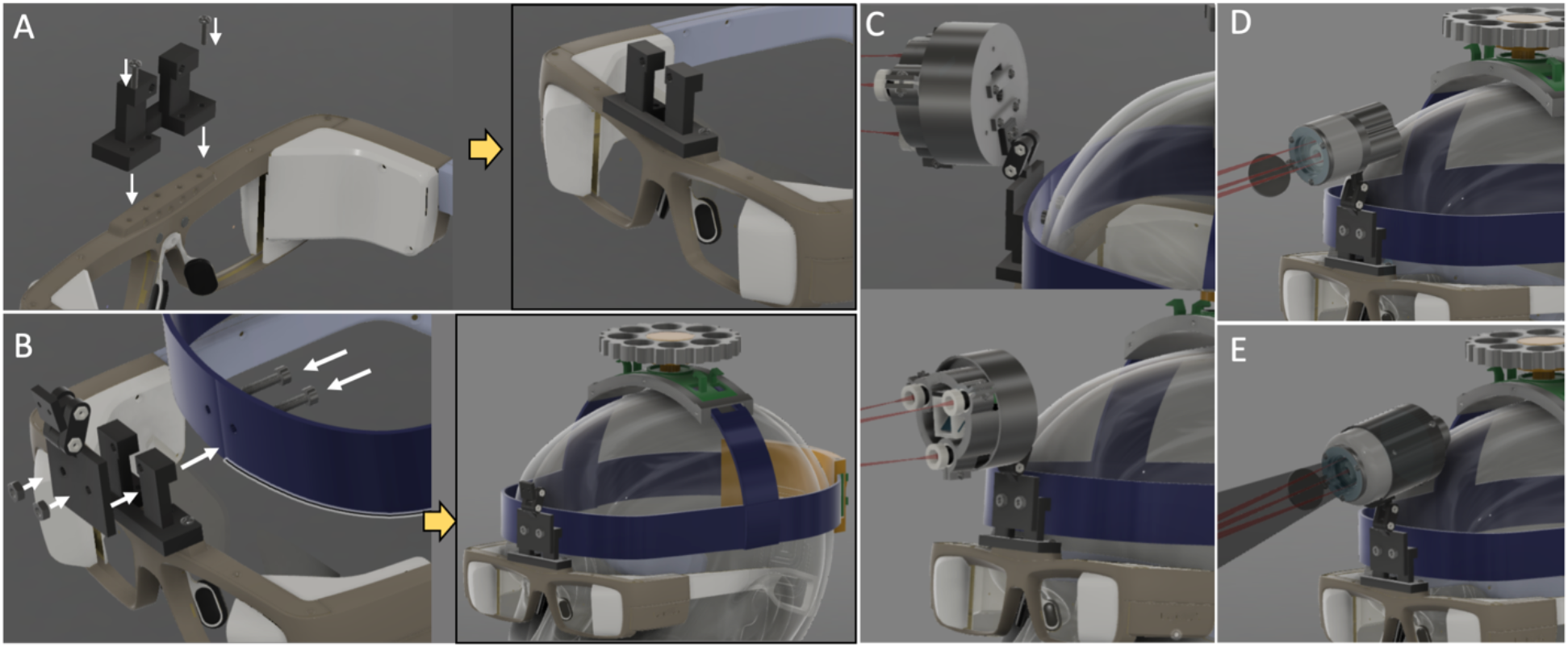
See-thru heads up display goggles can be mounted to the FAR-Pi surgical headband with a custom 3D printed adaptor. A) A 3D printed adaptor attaches to pre-existing mounting holes on the Lumus DK52 heads up display. B) The 3D printed adaptor has mounting holes that correspond to pre-existing mounting holes on the surgical headband forehead. A 3D printed articulating arm with the same mounting holes is secured together with the 3D printed adaptor. C) With the articulating mount in place, the illumination module with central imaging module is attached to the FAR-Pi system. D) and E) show alternative illumination modules (one with closely packed lasers and one with a central white light LED respectively) attached to the articulating mount.

**Fig. S13.**
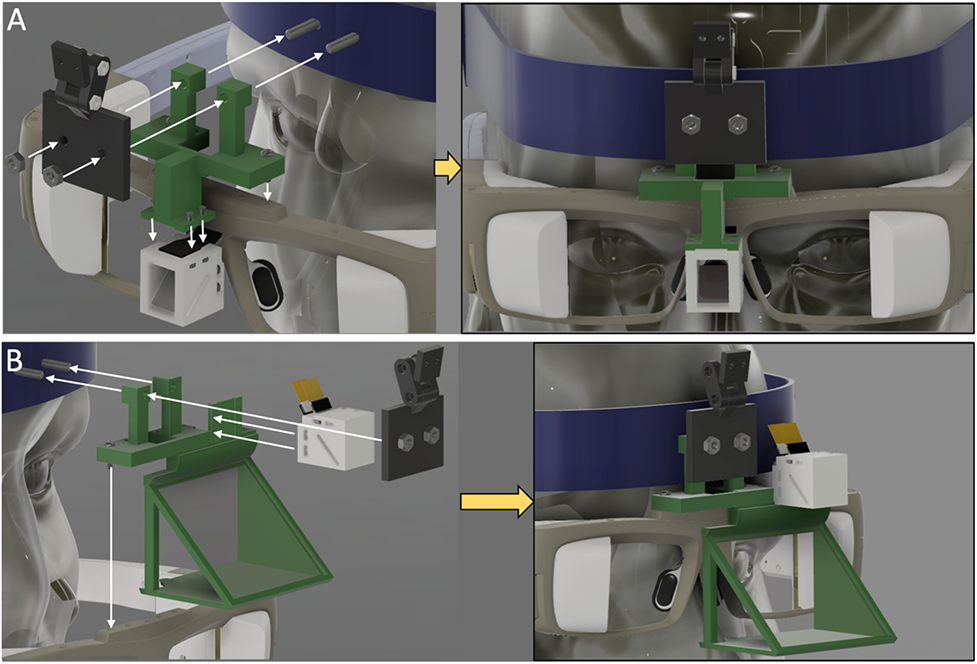
Assembly steps for mounting different implementations of the imaging module to the heads up display (HUD) glasses. A) To mount the imaging module between the eyes, the 3D printed adaptor that secures the HUD glasses and illumination module articulating arm to the surgical headband is adjusted to include M1.4 mounting holes that align with the back of the imaging module enclosure B) To make the imaging module coaxial with the eyes, a glass beamsplitter is employed. The 3D printed adaptor that secures the HUD glasses and illumination module articulating arm to the surgical headband is adjusted to include both a slot for a 50×50mm glass beamsplitter positioned at 45°, and a vertical mount with holes corresponding to the M1.4 mounting holes on the back of the imaging module enclosure.

#### 8. Fluorescent tissue phantom protocol

A 1 mM stock concentration of fluorophore (IR-125, 09030, Exciton Inc., Lockburne, OH)was achieved by dissolving 7.27 mg of IR-125 powder in 10 mL DMSO (Sigma Aldrich, St. Louis, MO). This concentration was validated using a spectrophotometer in which 1 µL of the 1 mM stock solution was dissolved in 999 µL of deionized water and calculated using the molar extinction coefficient of ICG in water. The ICG coefficient was used because 1) IR-125 has a nearly identical chemical structure and 2) the IR-125 molar extinction coefficient has only been characterized in methanol (232,000 cm^-1^ mol^-1^, from Exciton Inc. Certificate of Analysis) and was found to be comparable to ICG in methanol (221,000 cm^-1^ mol^-1^)(*50*). After concentration validation, the 1 mM IR-125 in DMSO stock solution was further diluted into 100 µM and 100 nm stocks with volumes of 1 mL each.

The base of the phantom was made using polyurethane (WC-783 A/B, BJB Enterprises, Tustin, CA) which consists of two parts (A and B) that are mixed together in a 10:9 part ratio. To begin, 20 mL of part A and 18 mL of part B were poured into a plastic disposable cup and mixed manually for 5 minutes. After mixing, 95 µL of the 100 nm stock was added and subsequently mixed to create an IR-125 concentration of 250 pM in polyurethane. Approximately 400 µL of the mixture was poured into two neighbor wells of the 96 well plate to achieve a 250 pM tissue mimicking phantom. Additional volumes of the IR-125 stock solution were serially added to the polyurethane mixture to create tissue phantom wells spanning 250 pM to 100 nM. Once all concentrations were poured into the 96 well plate, it was placed in a light-tight chamber at room temperature to cure overnight. Of note, when mixing the polyurethane and performing the serial dilutions, work quickly because curing begins in ∼30 minutes.

#### 9. Off the shelf component sourcing and cost

There are several different implementations of the FAR-Pi system, each implementation has the same set of core components with some implementation specific individual components. We list the parts that are needed to build the imaging module, illumination module, and computational module in tables S1-S3. The 3D printed parts that are used as enclosures and adaptors for each of the different FAR-Pi modules are not listed separately in the parts tables. In total these components require less than 250 grams of PLA, which can typically be purchased in 1 kg spools for $20. Assembly requires the use of a few dozen M1.4, M2 and M3 screws and bolts, which can be purchased in bulk kits containing hundreds of screws from Amazon.com for less than $20. Once assembled the imaging, illumination, and computational modules are mounted to a surgical headstrap. Adjustable surgical headbands can be purchased from Amazon.com, though they typically are packaged with additional components like a white light LED or a binocular loop which increases the total cost (Amazon.com ASIN: $65.99 #B01M3RF5GP, $56.99 #B0823QZSLL). We have found that the surgical headband alone can be 3D printed with PLA, and this makes purchase of a separate surgical headband unnecessary.

##### Imaging Module

Table S1 lists the components, cost, and vendor for the parts necessary to build the FAR-Pi imaging module. Because the imaging module can be built with different excitation filter options, the total cost of the imaging module is listed for each filter option.

While the FAR-Pi imaging module can be constructed for only $156 (option *a*), this option has limited fluorescence sensitivity and in-vivo SBR (see Figure 9, Main Text). The $200 filter (option *b*) however has excellent fluorescence sensitivity and in-vivo SBR, rivaling the more expensive filter *c* and *d* options.

##### Illumination Module

Table S2 lists the components, cost, and vendor for the parts necessary to build the FAR-Pi illumination module. We highlight 3 different designs, and for each design an additional excitation filter can be added. In design *a*, the imaging module is placed in the center. In design *b*, the imaging module is decoupled from the laser array and the lasers are closely packed without any space in the center. In design *c* there is space in the center of the lasers for a white light LED with IR cut off filter. For designs *a* and *c* individual clean-up filters are added to each laser diode. In design *b* the diodes are closely packed and a single clean-up filter can be utilized for all 3 laser diodes. The cost subtotals for each possible illumination module implementation are listed in Table S2.

##### Computational Module

Table S3 lists the components, cost, and vendor for the parts necessary to build the FAR-Pi computational module. The computational module has 2 different designs, one based on a Raspberry Pi 4B single board computer (RPiV4), and one with a Raspberry Pi Computational Module 4 (RPiCM4). The RPiV4 has only one camera CSI port, which is the input port that an RPiV2 camera requires. To support the use of two RPiV2 requires an additional Raspberry Pi Zero is used to convert one of the RPiV2 cameras into a USB camera that can be input to the RPiV4 via USB input. The RPiCM4 doesn’t have any output or input ports built in and requires an additional carrier board to exposure input and output. The additional cost of the RPiCM4 implementation is offset by the fact that the RPiCM4 supports two CSI input, obviating the need for an additional Raspberry Pi Zero. As a result the cost of both implementations is identical.

The total cost of the FAR-Pi system varies depending on choice of illumination module and imaging module filter. Table S4 summarize these costs for each unique implementation.

**Table S1.** Parts list for four implementations of the FAR-Pi imaging module showing component, quantity (#), total cost for the quantity of components listed, and vendor. Prices may be subject to change.

| **Component** | **#** | **Cost** | **Vendor** |
| --- | --- | --- | --- |
| RPiV2 standard (white light) camera module | 1 | \$ 25.00 | Raspberry Pi, multiple resellers |
| RPiV2 NOIR (NIR) camera module | 1 | \$ 30.00 | Raspberry Pi, multiple resellers |
| IMX219 Arducam sensor extension cables | 2 | \$ 25.98 | Arducam, #B0186 |
| IR Beamplitter - Cold IR Mirror | 1 | \$ 37.50 | Edmunds Optics, #62-634 |
| *Optical filter options (Options b-d showed significantly better performance than option a)* |  |  |  |
| a) Longpass filter - 830 nm CGA | 1 | \$ 38.00 | Newport, #5CGA-830 |
| b) Longpass filter - Double 830 nm CGA | 2 | \$ 76.00 | Newport, #5CGA-830 |
| c) Bandpass filter - OD 6 832 nm | 1 | \$ 225.00 | Edmund Optics, #84-091 |
| d) Longpass filter - 808 nm Edge Basic | 1 | \$ 525.00 | AVR Optics, BLP01-808R-12.5<br>*\$525 Price includes \$150 for custom sizing, waived if buying 10 or more |
| **Subtotals** |  |  |  |
| **Imaging module with filter option a** | **Imaging module with filter option b** | **Imaging module with filter option c** | **Imaging module with filter option d** |
| **\$ 156.48** | **\$ 194.48** | **\$ 343.48** | **\$ 643.48** |

**Table S2.**
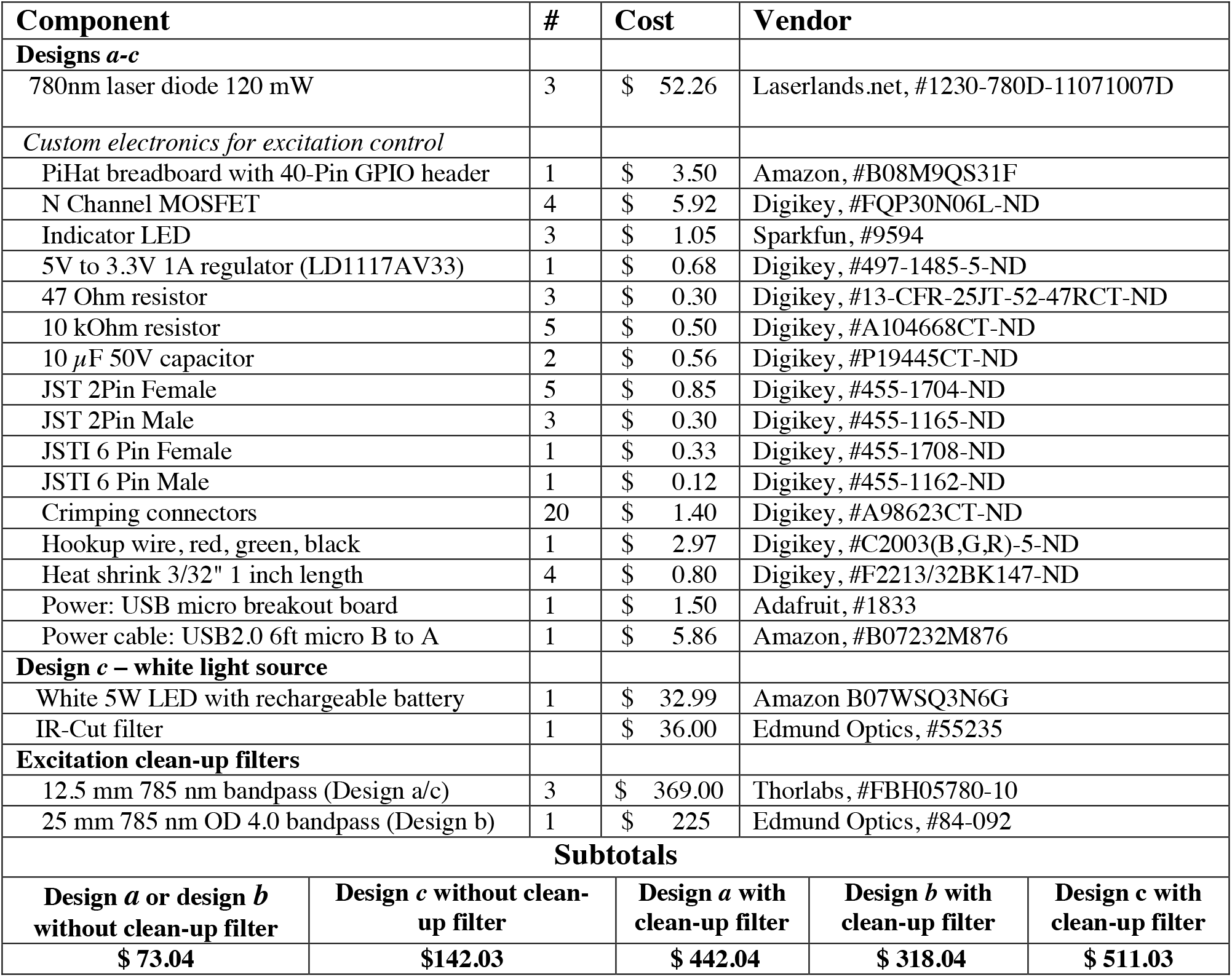
Parts and cost for 4 different implementations of the FAR-Pi illumination module with laser control circuit, showing component, quantity (#), total cost for the quantity of components listed, and vendor. Design *a* positions imaging module in the center of a laser diode array. Design *b* positions the laser diodes in close proximity. Design *c* positions a white light LED in the center of the laser array. Prices may be subject to change.

| Component | # | Cost | Vendor |  |  |
| --- | --- | --- | --- | --- | --- |
| Designs *a-c* |  |  |  |  |  |
| 780nm laser diode 120 mW | 3 | \$ 52.26 | Laserlands.net, #1230-780D-11071007D | | |
| Custom electronics for excitation control |  |  |  |  |  |
| PiHat breadboard with 40-Pin GPIO header | 1 | \$ 3.50 | Amazon, #B08M9QS31F | | |
| N Channel MOSFET | 4 | \$ 5.92 | Digikey, #FQP30N06L-ND | | |
| Indicator LED | 3 | \$ 1.05 | Sparkfun, #9594 | | |
| 5V to 3.3V 1A regulator (LD1117AV33) | 1 | \$ 0.68 | Digikey, #497-1485-5-ND | | |
| 47 Ohm resistor | 3 | \$ 0.30 | Digikey, #13-CFR-25JT-52-47RCT-ND | | |
| 10 kOhm resistor | 5 | \$ 0.50 | Digikey, #A104668CT-ND | | |
| 10 μF 50V capacitor | 2 | \$ 0.56 | Digikey, #P19445CT-ND | | |
| JST 2Pin Female | 5 | \$ 0.85 | Digikey, #455-1704-ND | | |
| JST 2Pin Male | 3 | \$ 0.30 | Digikey, #455-1165-ND | | |
| JSTI 6 Pin Female | 1 | \$ 0.33 | Digikey, #455-1708-ND | | |
| JSTI 6 Pin Male | 1 | \$ 0.12 | Digikey, #455-1162-ND | | |
| Crimping connectors | 20 | \$ 1.40 | Digikey, #A98623CT-ND | | |
| Hookup wire, red, green, black | 1 | \$ 2.97 | Digikey, #C2003(B,G,R)-5-ND | | |
| Heat shrink 3/32" 1 inch length | 4 | \$ 0.80 | Digikey, #F2213/32BK147-ND | | |
| Power: USB micro breakout board | 1 | \$ 1.50 | Adafruit, #1833 | | |
| Power cable: USB2.0 6ft micro B to A | 1 | \$ 5.86 | Amazon, #B07232M876 | | |
| Design *c* – white light source |  |  |  |  |  |
| White 5W LED with rechargeable battery | 1 | \$ 32.99 | Amazon B07WSQ3N6G | | |
| IR-Cut filter | 1 | \$ 36.00 | Edmund Optics, #55235 | | |
| Excitation clean-up filters |  |  |  |  |  |
| 12.5 mm 785 nm bandpass (Design *a/c*) | 3 | \$ 369.00 | Thorlabs, #FBH05780-10 | | |
| 25 mm 785 nm OD 4.0 bandpass (Design *b*) | 1 | \$ 225 | Edmund Optics, #84-092 | | |
| Subtotals |  |  |  |  |  |
| Design *a* or design *b* without clean-up filter | Design *c* without clean-up filter |  | Design *a* with clean-up filter | Design *b* with clean-up filter | Design *c* with clean-up filter |
| \$ 73.04 | \$142.03 | | \$ 442.04 | \$ 318.04 | \$ 511.03 |

**Table S3.** Parts and cost for 4 different implementations of the FAR-Pi computational module, showing component, quantity (#), total cost for the quantity of components listed, and vendor. Prices may be subject to change.

| **Component** | **#** | **Cost** | **Vendor** |
| --- | --- | --- | --- |
| **Raspberry PI V4 based (RPiV4)** |  |  |  |
| Computer - Raspberry Pi v4 4GB | 1 | \$ 55.00 | Raspberry Pi, multiple resellers |
| Sandisk 64GB Class 10 micro SD | 1 | \$ 12.19 | Amazon, #B08GYBBBBH |
| *Converting one RPiV2 camera to USB camera \*not needed if using Raspberry Pi Computer Module 4* |  |  |  |
| Raspberry Pi Zero | 1 | \$ 5.00 | Raspberry Pi, multiple resellers |
| CSI cable Pi Zero | 1 | \$ 5.95 | Adafruit, #3157 |
| Sandisk 32GB class 10 microSD | 1 | \$ 6.85 | Amazon #B010NE3QHQ |
| **Raspberry Pi CM4 based (RPiCM4)** |  |  |  |
| Computer- Raspberry PI Compute Module 4 4GB RAM 16 GB EEMC | 1 | \$ 65.00 | Seeed Studio, CM4102032 |
| Carrier board with two CSI inputs | 1 | \$ 19.99 | Waveshare, CM4-IO-BASE-A |
| **Components for either assembly** |  |  |  |
| Raspberry Pi USB Pi Switch | 1 | \$ 7.99 | Amazon, #B089ZVK2T6 |
| Cooling Fan, 5V 0.2A | 1 | \$ 2.95 | Digikey, #1528-4468-ND |
| USB Quick Charge A to C power cable 6ft | 2 | \$ 10.99 | Amazon, #B077Z2LCG8 |
| 30 cm micro HDMI to micro HDMI | 1 | \$ 11.59 | Amazon, #B019BVYAZO |
| Power bank - Anker 5V 4.8A 20,100mAh | 1 | \$ 59.99 | Amazon, #B00X5RV14Y |
| Power splitter | 1 | \$ 5.88 | Amazon, #B07CKQSTCB |
| **Subtotals** |  |  |  |
| RPiV4-based computational Module |  | RPiCM4-based computational module |  |
| **\$ 184.38** | | **\$ 184.38** | |

**Table S4.**
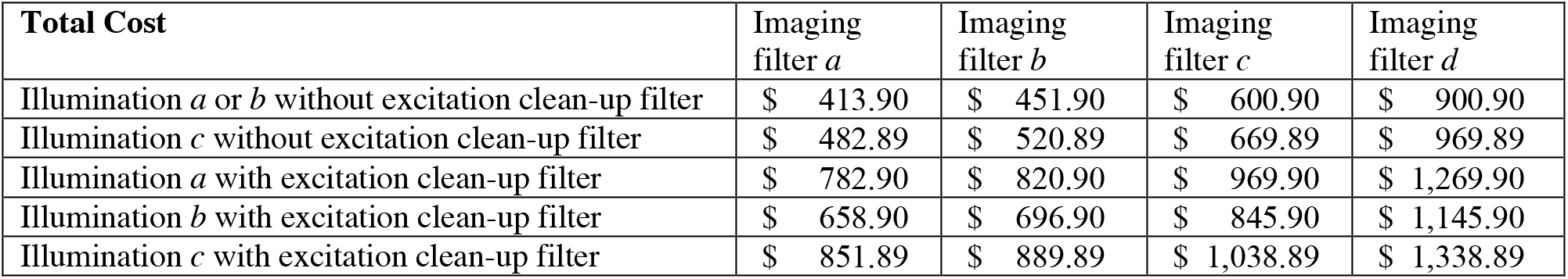
Total component cost of assembled FAR-Pi system depends on choice illumination and imaging modules.

